# CREB repressor in mushroom body enhances *Drosophila* LTM formation

**DOI:** 10.1101/2021.06.10.447902

**Authors:** Chun-Chao Chen, Hsuan-Wen Lin, Kuan-Lin Feng, Ruei-Yu Jhang, Linyi Chen, J. Steven de Belle, Tim Tully, Ann-Shyn Chiang

## Abstract

Long-term memory (LTM) requires learning-induced synthesis of new proteins allocated to specific neurons and synapses in a neural circuit. Not all learned information, however, becomes permanent memory. How the brain gates relevant information into LTM remains unclear. In *Drosophila* adults, a single training session in an olfactory aversive task is not sufficient to induce protein synthesis-dependent LTM. Instead, multiple spaced training sessions are required. Here, we report that initial learning induces neural activity in the early α/β subset of Kenyon cells of the mushroom body (MB), and output from these neurons inhibits LTM formation. Specifically in response to spaced training, Schnurri activates CREBB expression which then appears to suppress the inhibitory output from MB. One training session can enhance LTM formation when this inhibitory effect is relieved. We propose that learning-induced protein synthesis and spaced training-induced CREBB act antagonistically to modulate output from early α/β MB neurons during LTM formation.

## Introduction

*Drosophila* continues to demonstrate its utility as a model system to study memory, more than four decades after the first mutant was described (Dudai et al., 1976). Genetic dissection of olfactory aversive memory formation using various single-gene mutants has revealed at the behavioral level several distinct temporal phases, including short-term memory (STM), middle-term memory (MTM), anesthesia-resistant memory (ARM) and long-term memory (LTM) (Tully et al., 1994; Tully et al., 1990; Tully, 1996; Quinn and Dudai, 1976). The initial learning event (acquisition) after a single training session (1x) appears to induce STM, MTM and ARM, while spaced training (10 training sessions with 15 min rest intervals between each, 10xS) appears uniquely required to induce LTM consolidation. Manipulations of several of these “memory genes” also have established cases where memory formation is either impaired or enhanced, revealing bi-directional biochemical modulation of memory formation (Yin et al., 1994; Yin et al., 1995a; Ge et al., 2004; Presente et al., 2004; Wu et al., 2007; Pavlopoulos et al., 2008; Huang et al., 2012; Tubon et al., 2013; Fropf et al., 2014; Lee et al., 2018; Scheunemann et al., 2018).

As the neural substrates of olfactory memory formation are elucidated in flies, a remarkable “memory circuit” is emerging. Olfactory information delivered from the antennal lobe (AL) by projection neurons (PN) and foot shock reinforcement delivered by dopaminergic neurons (DAN) both converge on mushroom body (MB) neurons in the central brain where their coincidence triggers cascading cellular events that underlie learning (Dubnau and Chiang 2013; Perisse et al., 2013; Davis, 2015; Cognigni et al., 2018). MBs play a predominant role in subsequent memory formation, together with several groups of extrinsic MB neurons. Sequential genetically-defined memory phases map onto distinct subpopulations of these neurons. STM involves γ, α′/β′ and α/β neurons and two classes of MB output neurons (MBON: MB-M4, MB-M6) (Blum et al., 2009; Scheunemann et al., 2012; Bouzaiane et al., 2015). MTM involves neural activity in γ, α/β and MB-V2 neurons (Blum et al., 2009; Scheunemann et al., 2012; Bouzaiane et al., 2015). ARM requires neural activity in MB γ, α′/β′, α/β neurons, dorsal paired medial (DPM) neurons, anterior paired lateral (APL) neurons, DAN and four different MB output neurons (MB-M4, MB-M6, MB-V2, MBON-β2β′2a) (Lee et al., 2011; Knapek et al., 2011; Placais et al., 2012; Wu et al., 2013; Bouzaiane et al., 2015; Yang et al., 2016; Scholz-Kornehl and Schwärzel, 2016; Kotoula et al., 2017; Shyu et al., 2019). LTM involves neural activity in late MB α/β neurons with output from DPM, serotonergic projection neurons (SPN) and three classes of MBONs (MB-V3, MB-M4, MBON-γ3,γ3β′1). Cyclic AMP response element binding protein (CREB)-dependent consolidation of LTM also requires activity in dorsal anterior lateral (DAL) neurons (Chen et al., 2012; Pai et al., 2013; Tonoki and Davis 2015; Bouzaiane et al., 2015; Wu et al., 2017; Scheunemann et al., 2018). Finally, memory retrieval depends on neural activity in DAL, pioneer α/β neurons and four classes of MBONs (MB-V2, MB-V3, MB-M4, MBON-γ3,γ3β′1) (Séjourné et al., 2011; Chen et al., 2012; Pai et al., 2013; Bouzaiane et al., 2015; Wu et al., 2017).

Here, we describe another enlightening property of olfactory memory in *Drosophila*: inhibition of LTM formation at the circuit level. Output from the early α/β subpopulation of MB neurons appears initially to inhibit LTM formation, but with spaced training, transcription of *crebB* (*dcreb2*, repressor) is induced therein, apparently reducing neural output therefrom and thereby enabling LTM formation. Thus, persistent olfactory memory formation appears modulated or “gated” at the level of neural activity in early α/β MB neurons. These observations presage the need for a more general deconvolution of biochemical mechanisms into distinct neuronal subtypes within a memory circuit.

## Results

### Learning inhibits LTM

Sixty of the single-gene mutants mentioned above were generated using transposon mutagenesis and were screened for impairments of memory one day after 10xS training (Dubnau et al., 2003). Twenty-two of these lines carried *P-Gal4* enhancer traps, which enabled us to drive targeted inducible expression of a temperature-sensitive *Ricin^CS^* transgene and then block protein synthesis after 10xS (Chen et al., 2012; Pai et al., 2013; Wu et al., 2017). With protein synthesis inhibited in this manner, we found impairments of 1-day memory in nine of these lines (figure supplement 1A-C; table supplement 1) (Tully et al., 1994; Yin et al., 1994). Remarkably, GFP was expressed in DAL neurons in all nine cases (figure supplement 1B), an observation that contributed to our characterization of DAL neurons extrinsic to the MB as *bona fide* “LTM neurons” (Chen et al., 2012). Two of the enhancer-trap memory mutants that we screened were particularly informative. In *umnitza* flies, GFP was expressed in DAL neurons but very weakly in MB, and 1-day memory after 10xS was impaired. Conversely in *norka* flies, GFP was expressed in MB but not in DAL neurons and 1-day memory after 10xS was normal.

What then might be going on in the seven enhancer-trap mutants with normal memory and with GFP expression in both MB & DAL neurons (Figure 1A; figure supplement 1A)? First, we confirmed that active Ricin^CS^ inhibition of protein synthesis in different subsets of MB neurons did not impair 1-day memory after 10xS (figure supplement 2A-B and 3). In contrast, 1-day memory after 10xS was impaired by active Ricin^CS^ in DAL or MB-V3 neurons (Chen et al., 2012; Pai et al., 2013; Wu et al., 2017) but was normal after massed training (10x training sessions with no rest intervals; 10xM) or in control flies with inactive Ricin^CS^ (18 °C) (figure supplement 3).

**Figure 1.**
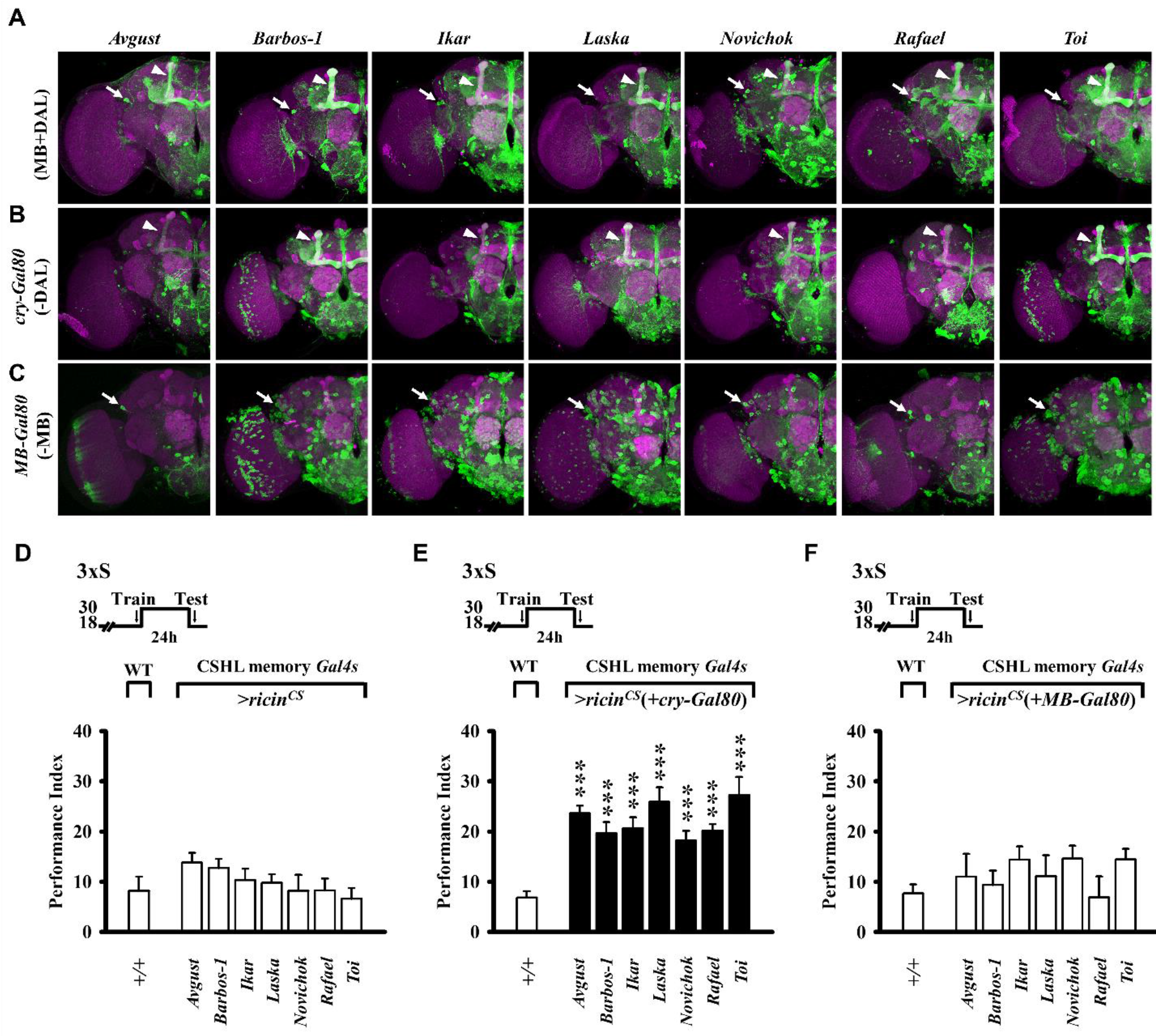
CSHL memory *Gal4* patterns in which protein synthesis inhibition has no net effect on LTM formation. (**A**) Full expression patterns that include MB and DAL neurons (top). (**B**) Expression restricted from DAL neurons by *cry-Gal80* inhibition of Gal4 (center). (**C**) Expression restricted from MB neurons by *MB-Gal80* inhibition of Gal4 (bottom). DAL (arrow) and MB (arrowhead). (**D-F**) Protein synthesis inhibition in MB neurons enhances LTM formation. Effects of memory circuit *Gal4*-targeted Ricin^CS^ on 1-day memory after three spaced training cycles (3xS), compared with wild-type (+/+) controls. Cold-sensitive Ricin^CS^ blocks protein synthesis at the permissive temperature (30 °C) between training and testing. (**D**) Protein synthesis inhibition in all Gal4-expressing elements of the memory circuit has no net effect on LTM. (**E**) Ricin^CS^ expression and blocking of protein synthesis in MB but not in DAL neurons (where *cry-Gal80* inhibits Gal4), enhances 1-day memory in all seven *Gal4* genotypes. (**F**) By contrast, Ricin^CS^ expression and blocking of protein synthesis in DAL but not in MB neurons (where *MB-Gal80* inhibits Gal4) has no effect on LTM. In all figures, temperature control schedules are indicated (top). Memory performance indices are calculated as the normalised percent avoidance of shock-paired odour. Bars represent mean ± SE, *n* = 8/bar unless stated otherwise. *, *P* < 0.05; **, *P* < 0.01, ***, *P* < 0.001. All genotypes are listed in table supplement 2.

We next tested the hypothesis that inhibition of protein synthesis in MB might enhance LTM formation, thereby off-setting the impairment produced by blocking protein synthesis in DAL neurons. Using *cry-Gal80* or *MB-Gal80*, we blocked transgenic Ricin^CS^ expression outside (i.e. DAL neurons) or inside of MB, respectively, in these seven enhancer-trap memory lines (Figure 1B-C) and then subjected them to suboptimal 3xS training. Surprisingly, LTM formation was *enhanced* in all seven lines by *cry-Gal80* subtraction (Figure 1D-F).

The MB is composed of approximately 2,500 intrinsic neurons (Kenyon cells; KCs) developmentally derived from four neuroblasts and distinguished by their projections that form the γ, α′/β′ and α/β lobes (Ito et al., 1997; Zhu et al., 2003; Lin et al., 2007). We looked among these neuronal subpopulations to identify where LTM enhancement might reside (Figure 2; figure supplement 2A). Active Ricin was expressed in all KCs or in γ, α′/β′, or α/β neurons separately. Enhanced LTM was observed after 3xS only in α/β neurons, which were previously shown to have a role in LTM formation (Blum et al., 2009; Yu et al., 2006) (Figure 2A).

**Figure 2.**
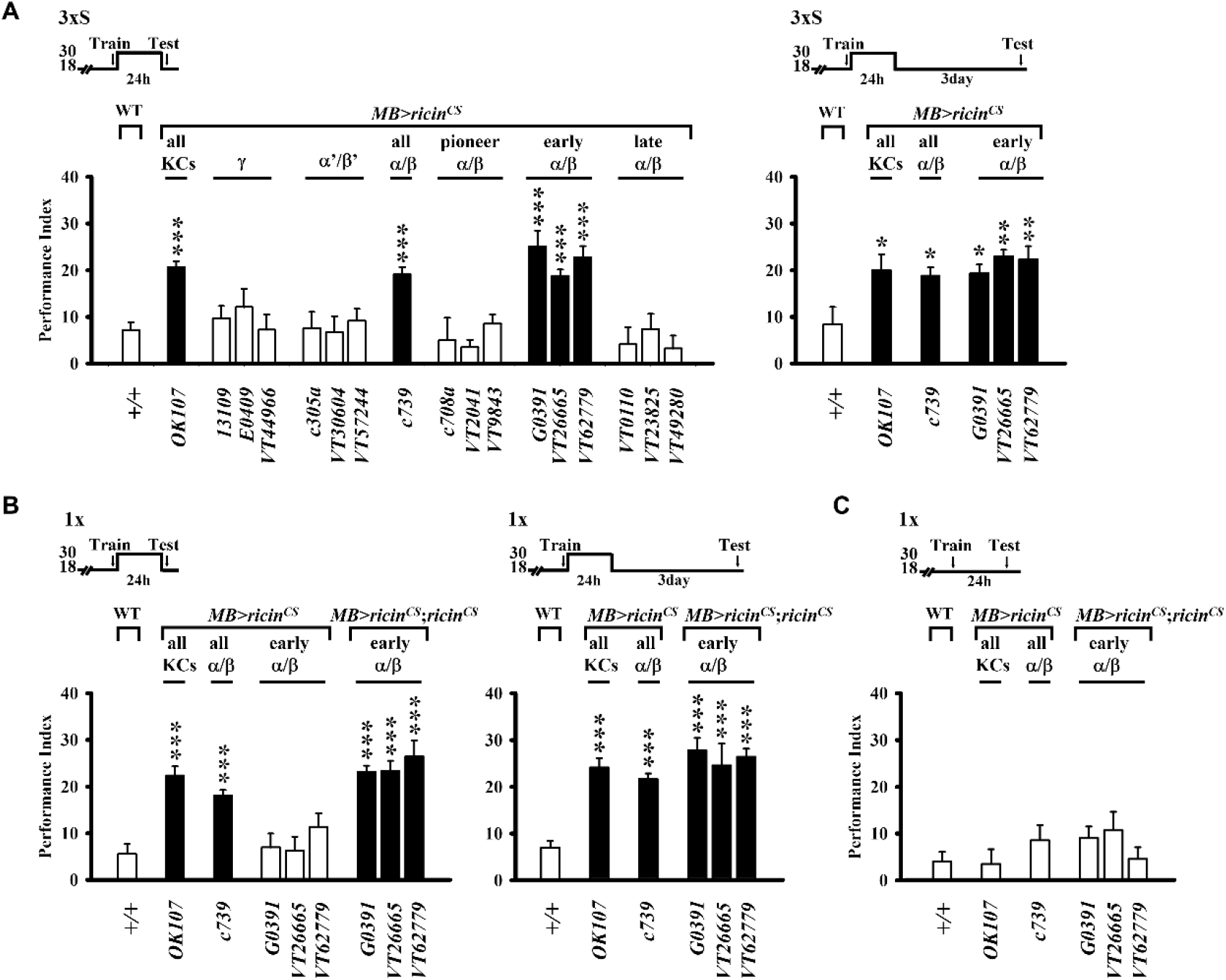
Protein synthesis inhibition in early α/β MB neurons enhances LTM formation. Effects of MB *Gal4*-targeted Ricin^CS^ on memory after sub-threshold training compared with wild-type controls (see MB *Gal4* expression patterns in figure supplement 2A) (**A**) Protein synthesis inhibition in MB *Gal4* patterns that include early α/β neurons enhance 1-day (left) and 4-day (right) memory after 3xS training. (**B**) Two copies of Ricin^CS^ expressed in early α/β neurons are necessary to enhance 1-day (left) and 4-day (right) memory after 1x training, whereas only one copy of Ricin^CS^ is insufficient. (**C**) Expression of inactive Ricin^CS^ in α/β neurons at the restrictive temperature (18 °C) has no effect on memory after 1x training.

The α/β neurons are subdivided further into three types: pioneer α/β, early α/β and late α/β neurons based on their birth sequences (Zhu et al., 2003; Lin et al., 2007; Tanaka et al., 2008; Aso et al., 2014). When active Ricin was expressed in these three subpopulations separately, we observed enhanced LTM after 3xS only when protein synthesis was blocked in early α/β neurons (Figure 2A, left). Enhanced LTM lasted for at least 4 days (Figure 2A, right) and was not observed in control flies (inactive Ricin^CS^) after 3xS training or in flies with active Ricin^CS^ after 3xM (figure supplement 2C). Blocking protein synthesis in early α/β neurons enhanced 1- and 4-day memories even after only 1x training (Figure 2B). Notably, LTM enhancement after 1x training required two copies of transgenic Ricin^CS^. Together, these results indicate that inhibition of protein synthesis in early α/β neurons yields a *bona fide* enhancement of LTM formation.

We next inquired about the most effective time after training when inhibition of protein synthesis would enhance LTM. Ricin^CS^ in early α/β neurons was activated for 12 h in a series of time windows staggered by 2 h during the first 24 h after 1x training (Chen et al., 2012; Wu et al., 2017). LTM was enhanced when protein synthesis was blocked beginning from 0- to 4-h but not from 6- to 12-h after training (Figure 3A). Shortening the inhibition period to 3 h, we resolved the window of protein-synthesis-dependent LTM inhibition to the first 6 h after training (Figure 3B).

**Figure 3.**
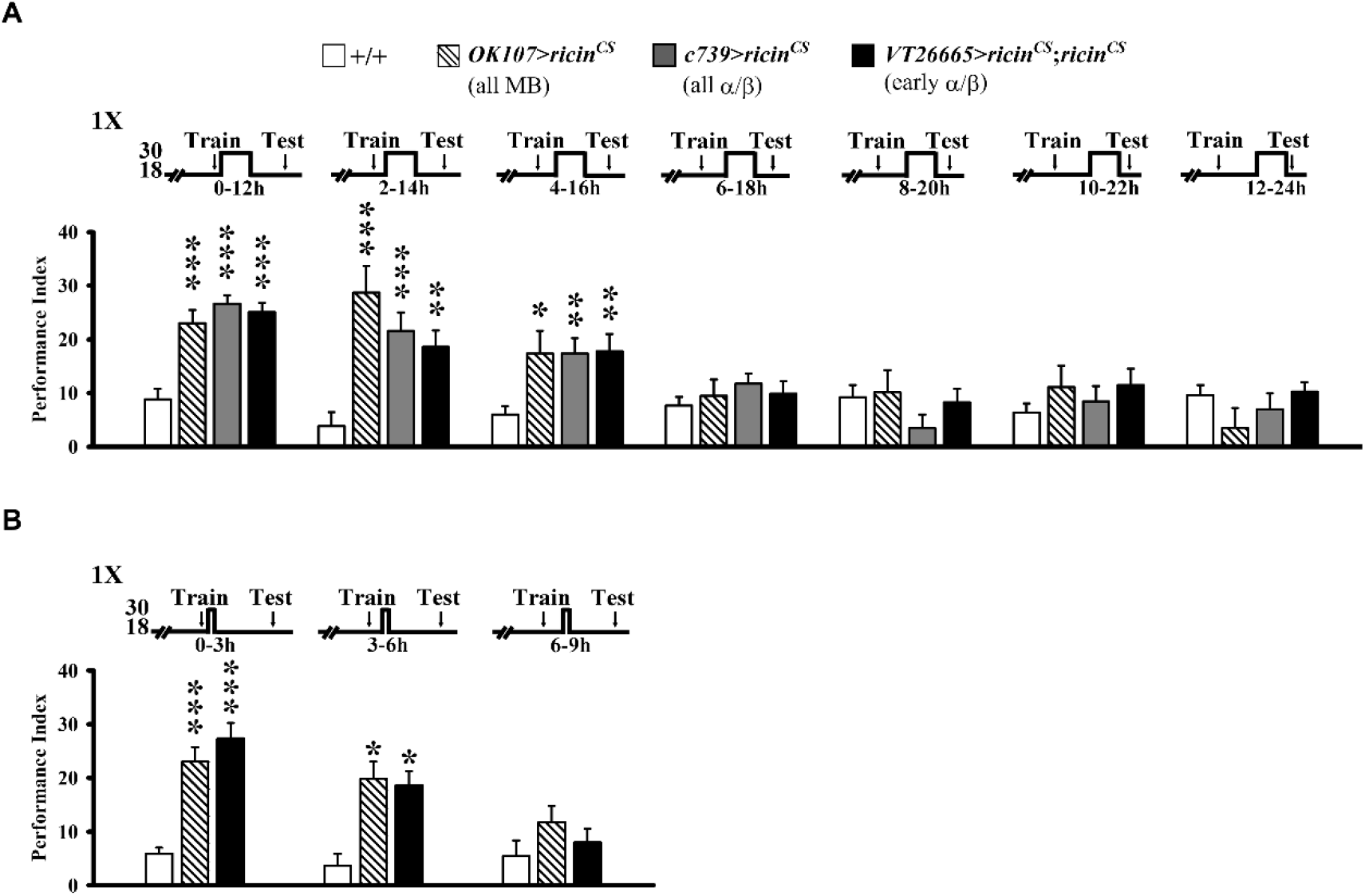
Protein synthesis in early α/β neurons after subthreshold training antagonizes LTM formation. Blocking protein synthesis in all MB (OK107), all α/β (c739), and early α/β (VT26665) neurons 0-6 h after 1x training enhances 1-day memory compared with wild-type (+/+) controls (related to Figure 2; also compare with Figure 4A). (**A**) Memory enhancement is significant in all groups with Ricin^CS^ activity 0-12 h, 2-14 h and 4-16 h after training but not for later time windows. (**B**) Memory enhancement is significant in all groups with Ricin^CS^ activity 0-3 h and 3-6 h but not 6-9 h after training.

### Output from early α/β neurons inhibits LTM

To address whether the inhibitory effect on LTM in early α/β lobes depends on their neural output, we blocked synaptic transmission from pioneer α/β, late α/β or early α/β neurons using *UAS-shi^ts^* (Dubnau et al., 2001; McGuire et al., 2001). In these experiments we found that 1- and 4-day memory after 3xS training were enhanced by this manipulation in early α/β (Figure 4A-B), but not in pioneer or in late α/β neurons (figure supplement 4A). Interestingly, we also established that LTM after 3xS was enhanced when synaptic transmission was blocked from early α/β during the first 8 h period after training but not 9-24 h after training or during 1-day memory retrieval (Figure 4A). Thus, this temporal requirement for synaptic transmission from early α/β neurons corresponds to the requirement for protein synthesis (Figure 3). We confirmed these results using two additional *Gal4* drivers expressing specifically in early α/β neurons (Figure 4C; figure supplement 2A) and observed normal 1-day memory after 3xM (Figure 4B, middle) as in control flies carrying *UAS-shi^ts^* alone, *Gal4* alone (Figure 4B, right) or at the permissive temperature for *shi^ts^* (Figure 4A, left and 4C, right). Finally, we established that LTM after 10xS training was not impaired when neural output from early α/β neurons was blocked (figure supplement 4B). Together, these results indicate that, in the absence of spaced training, neural output from early α/β neurons inhibits LTM formation. To examine the influence of early α/β neuron membrane excitability on memory, we used ectopic expression of either transgenic hyperexcitors (*UAS-Shaw^DN^*, a dominant-negative Shaw potassium channel and *UAS-NaChBac*, a sodium channel) or hypoexcitors (*UAS-Shaw*, a Shaw potassium channel and *UAS-Kir2.1::GFP*, an inward-rectifying potassium channel Venken et al., 2011). Temporal control of these transgenes was enabled using a *tub-Gal80^ts^* transgene (conditional expression of Gal80 suppresses Gal4 expression at 18 °C but not at 30°C McGuire et al., 2003). We found that increasing membrane excitability of early α/β neurons impaired 1-day memory after 10xS training (Figure 5A, left) without affecting (1) memory after 1x training (fig. S5A), (2) 1-day memory after 10xM training (figure supplement 5B) or (3) 1-day memory after 10xS training when flies were kept at permissive temperature (18°C) (figure supplement 5C). This inhibitory effect appears to be complete, because inhibition of protein synthesis by feeding flies cycloheximide (CXM) did not further reduce 1-day LTM after 10xS training (Figure 5A, right). Decreasing membrane excitability of early α/β neurons with ectopic expression of hypoexciter transgenes, on the other hand, enhanced both 1- and 4-day memory after 1x training (Figure 5B, left and middle), whereas enhancement of 1-day memory was not observed in control transgenic flies kept at the permissive temperature (18°C) (Figure 5B, right). We also found normal 1-day memory in these transgenic flies after 10xS training (Figure 5C). Thus, sufficient spaced training appeared to occlude the enhancing effects on LTM formation of decreased membrane excitability in early α/β neurons. These data further support the notion that neural activity from early α/β neurons inhibits LTM formation downstream (Dubnau and Chiang 2013; Pai, et al., 2013; Wu et al., 2017).

**Figure 4.**
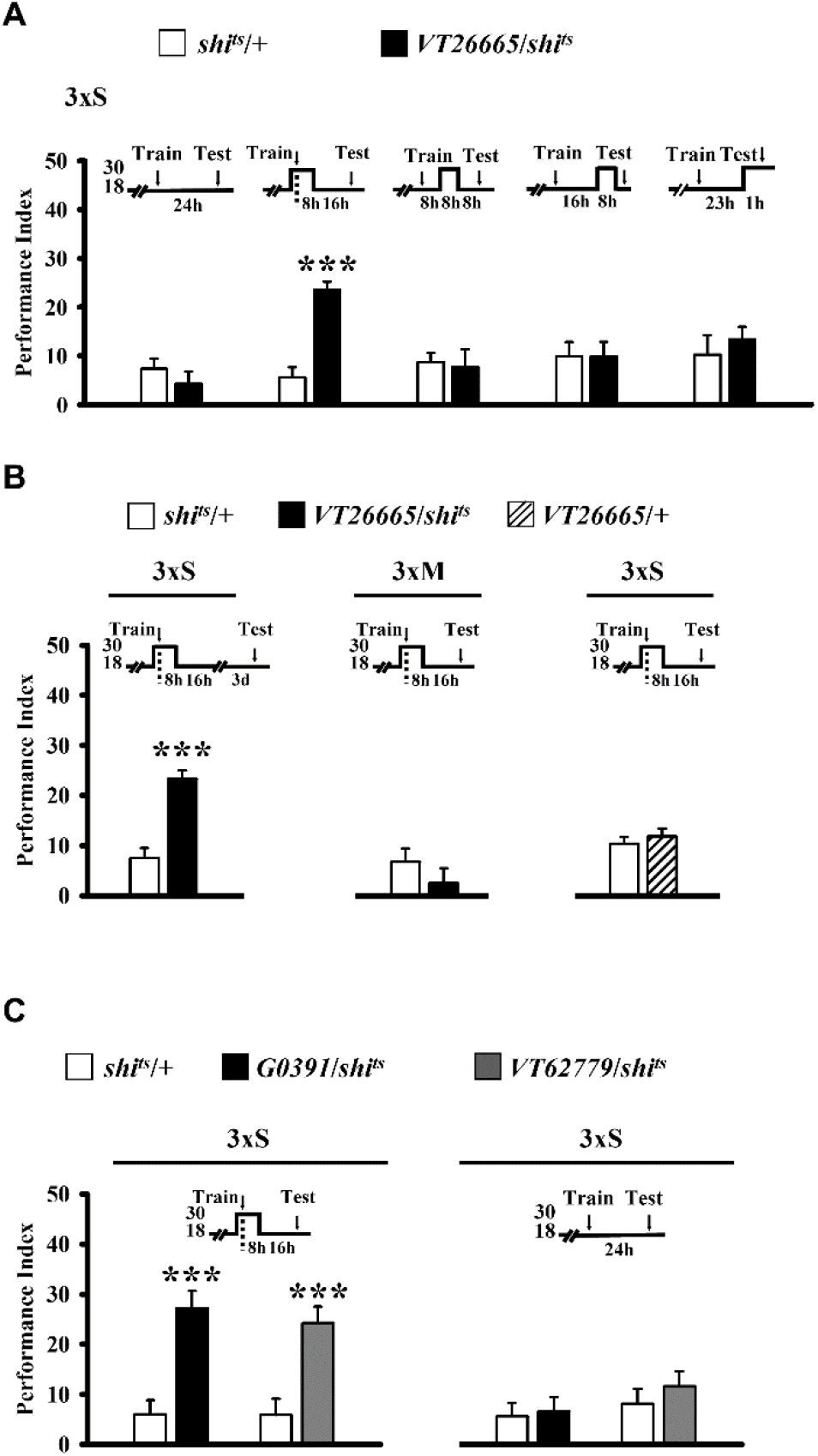
Blocking early a/b neuron signaling enhances LTM formation. Effects of early a/b *Gal4*-targeted Dynamin^ts^ (shi^ts^) on 1-day memory after 3xS training. (**A**) Blocking signaling in the first 8-h after training enhances 1-day memory, whereas blocking signaling during subsequent 8-h windows has no effect, compared with the unexpressed *shi^ts^/+* control (also see figure supplement 4). (**B**) Similarly, blocking signaling in the first 8-h after 3xS training enhances 4-day memory (left), but not after 3xM training (center) or after 3xS training of the *Gal4* driver or *shi^ts^* transgene alone (right). (**C**) Blocking early a/b signaling using two additional *Gal4* patterns confirms this inhibitory effect and enhancement of 1-day memory (left). Memory is unaffected in flies held at the permissive temperature (18 °C) after 3xS training (right).

**Figure 5.**
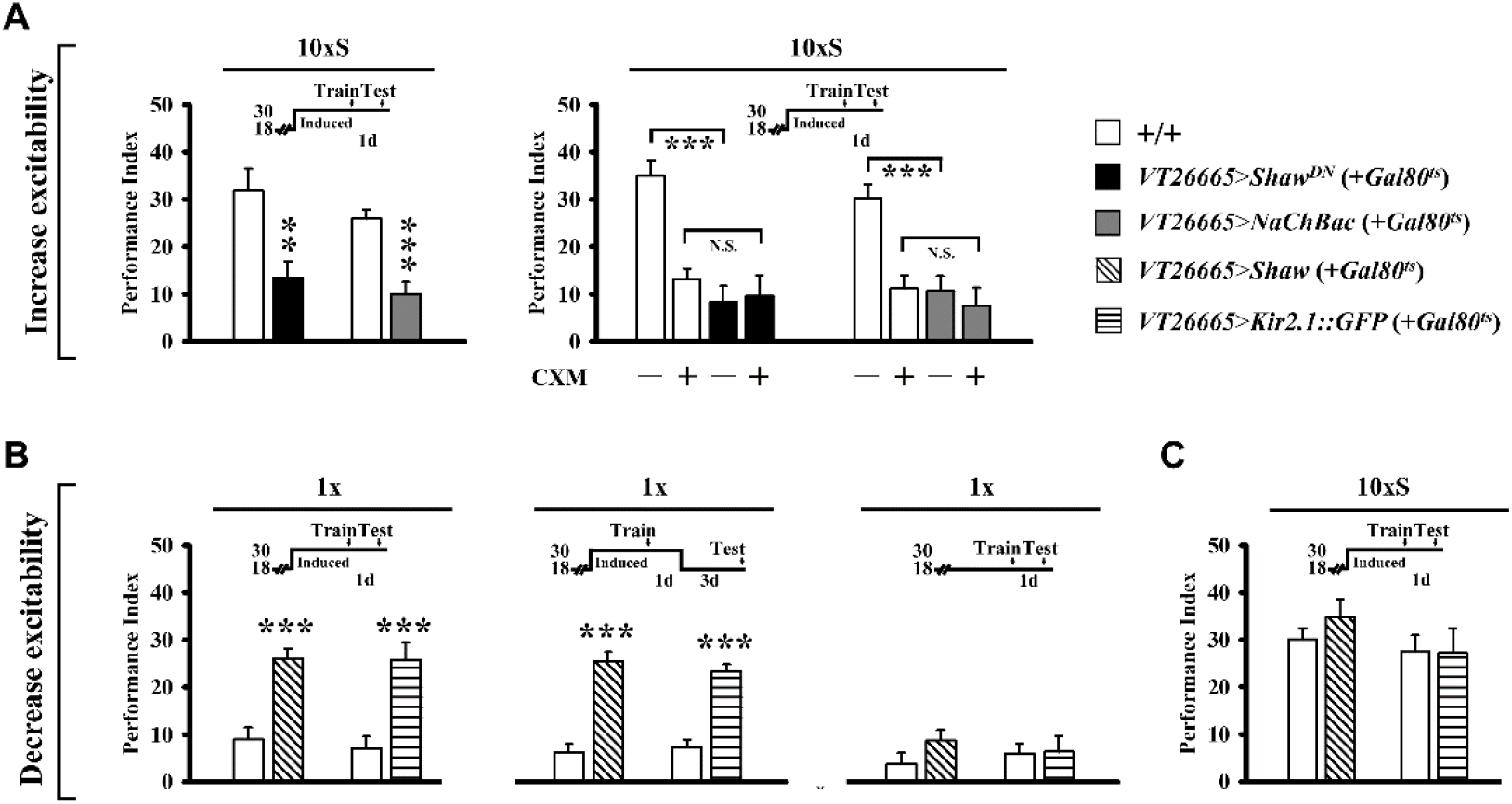
Neural membrane excitability in early α/β neurons bi-directionally regulates LTM formation. MB early a/b *Gal4*-targeted expression of K^+^ and Na^+^ channel proteins is induced at the restrictive temperature for the *tub-Gal80^ts^* inhibitor (30 °C) from five days before training until testing. (**A**) *Shaw^DN^* and *NaChBac* overexpression increase neural activity and impair 1-day memory after 10xS training (left). The same LTM is similarly blocked by systemic protein synthesis inhibition induced by CXM feeding (right) (also see figure supplement 5). (**B**) In contrast, *Shaw* and *Kir2.1::GFP* overexpression decrease neural activity and enhance1-day memory after only 1x training (left), which endures for at least 4 days (center). Memory is unaffected in these flies held at the permissive temperature for *tub-Gal80^ts^* (18 °C) after 1x training (right). (**C**) Decreasing neural activity as in (**B**) does not affect 1-day memory after 10xS training.

### cAMP signaling in early α/β neurons enhances LTM

Neural excitability is suggested to be modulated by cAMP signaling (Davis et al., 1998; Baines, 2003), which in MB is also involved in LTM formation (Blum et al., 2009). Accordingly, we inducibly overexpressed *rutabaga^+^* (rut^+^) adenylyl cyclase (AC) or constitutively active *cAMP-dependent protein kinase* (Pka^act1^) transgenes in early α/β neurons and found that 1-day memory after 1x training was enhanced to levels normally seen after 10xS in both cases (Figure 6A and 7A). Moreover, inducible RNAi knockdowns of these genes impaired 1-day memory after 10xS (Figure 6b and 7b; figure supplement 6). Together, these results suggest that LTM formation is also modulated by cAMP in early α/β neurons.

**Figure 6.**
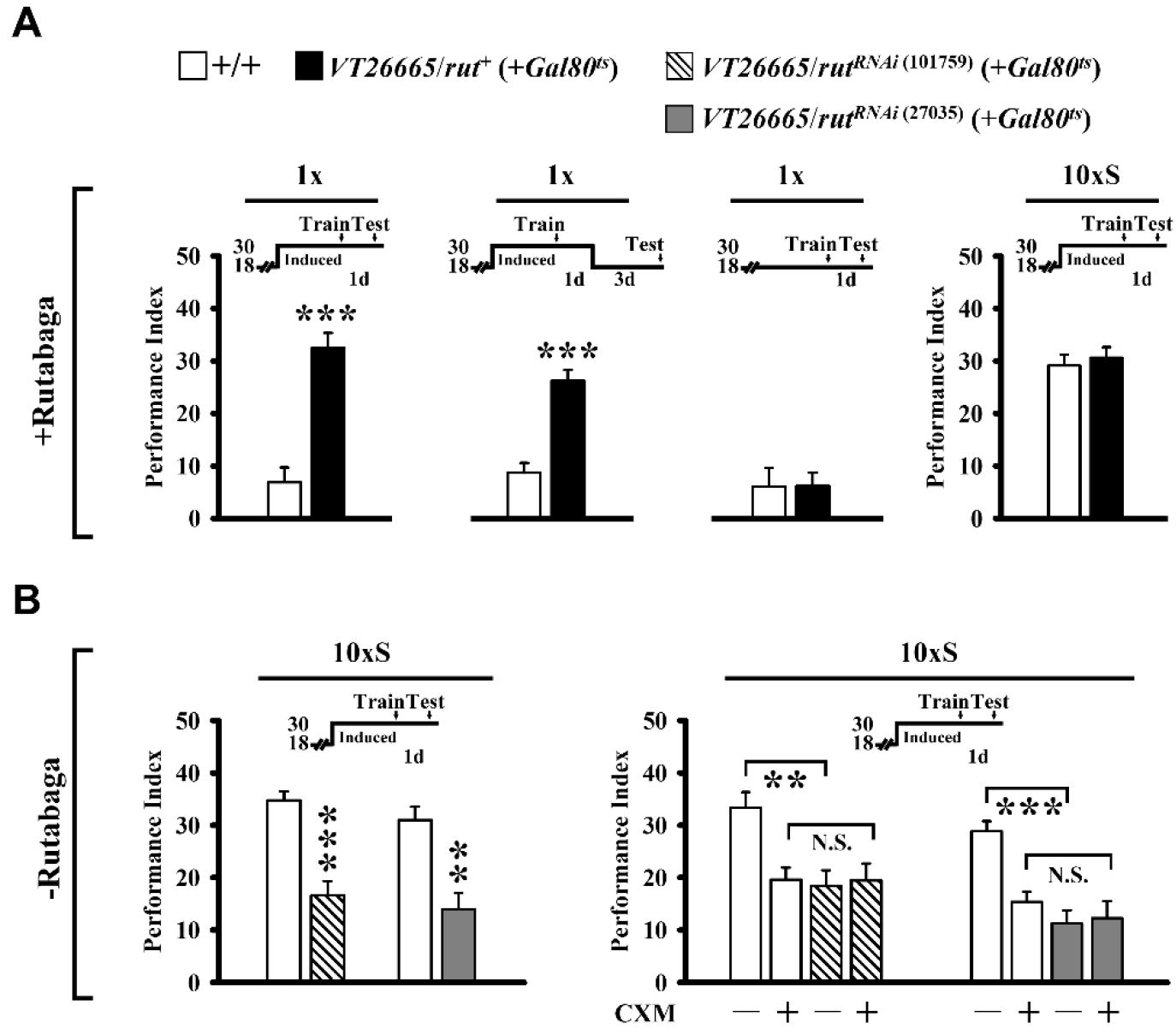
Modulation of Rutabaga (AC) in early α/β neurons bi-directionally regulates LTM formation. (**A**) Overexpressing AC in early α/β neurons enhances 1-day memory after only 1x training (left) and lasts at least four days (left center). Memory is unaffected in these flies held at the permissive temperature for *tub-Gal80^ts^* (18 °C) after 1x training (right center). One-day memory is unaffected in these flies held at 30 °C after 10xS training (right). *Gal4*-targeted *rut*^+^ overexpression is induced at the restrictive temperature for *tub-Gal80^ts^* (30 °C) from five days prior to training until testing. (**B**) By contrast, adult-stage specific RNAi down-regulation of AC in early α/β neurons *(two independent RNAi lines) impairs* 1-day memory after 10xS training (left) (also see figure supplement 6). The same LTM is similarly blocked by systemic protein synthesis inhibition induced by cycloheximide (CXM) feeding (right).

**Figure 7.**
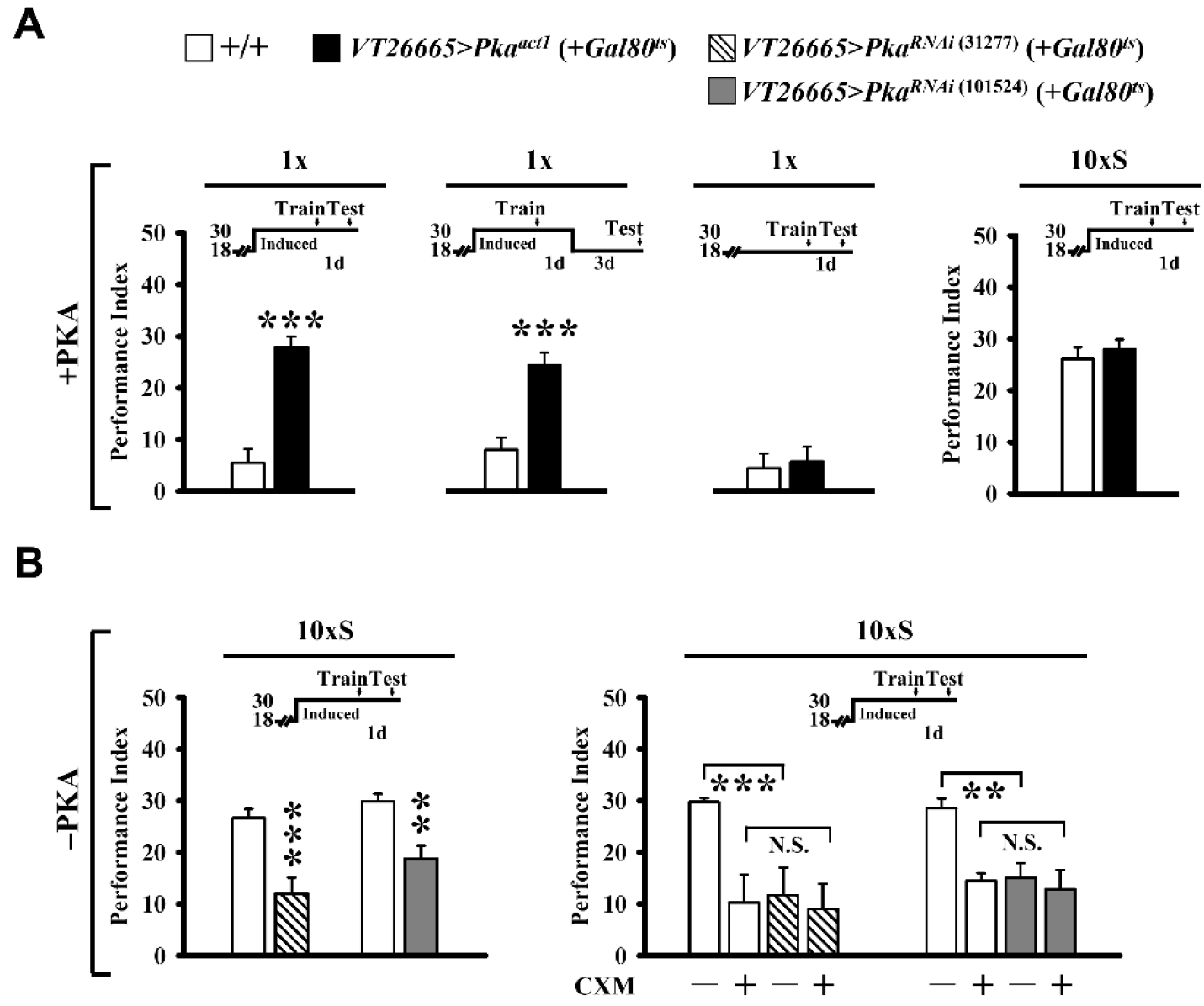
Modulation of PKA in early α/β neurons bi-directionally regulates LTM formation. (**A**) Overexpressing constitutively active PKA^act1^ in early α/β neurons enhances 1-day memory after only 1x training (left) and lasts at least four days (left center). Memory is unaffected in these flies held at the permissive temperature for *tub-Gal80^ts^* (18 °C) after 1x training (right center). One-day memory is unaffected in these flies held at 30 °C after 10xS training (right). *Gal4*-targeted PKA^act1^ overexpression is induced at the restrictive temperature for *tub-Gal80^ts^* (30 °C) from five days prior to training until testing. (**B**) By contrast, adult-stage specific RNAi down-regulation of PKA in early α/β neurons *(two independent RNAi lines) impairs* 1-day memory after 10xS training (left) (also see figure supplement 6). The same LTM is similarly blocked by systemic protein synthesis inhibition induced by cycloheximide (CXM) feeding (right).

### CREBB in early α/β neurons enhances LTM

Protein synthesis-dependent LTM formation also depends on CREBB transcription factors, and expression of CREBB protein is thought to be dependent on the expression level of protein kinases involved in cAMP signaling (Lee et al., 2018). Consistent with our Rutabaga and PKA knockdown results, we induced RNAi knockdown of CREBB in early α/β neurons and observed impairment of 1-day memory after 10xS training. Further impairment was not seen in combination with systemic protein synthesis inhibition by feeding CXM (Figure 8C). Because inhibition of protein synthesis in early α/β neurons instead produced an enhancing effect on LTM formation, we inducibly expressed *crebB* repressor transgenes (Zhang et al., 2019) in these neurons only, with the expectation that the manipulations would lead to enhanced memory. Indeed, expressing *crebB* enhanced 1-day memory after 1x training to levels normally seen after 10xS training (Figure 8A-B; figure supplement 6).

**Figure 8.**
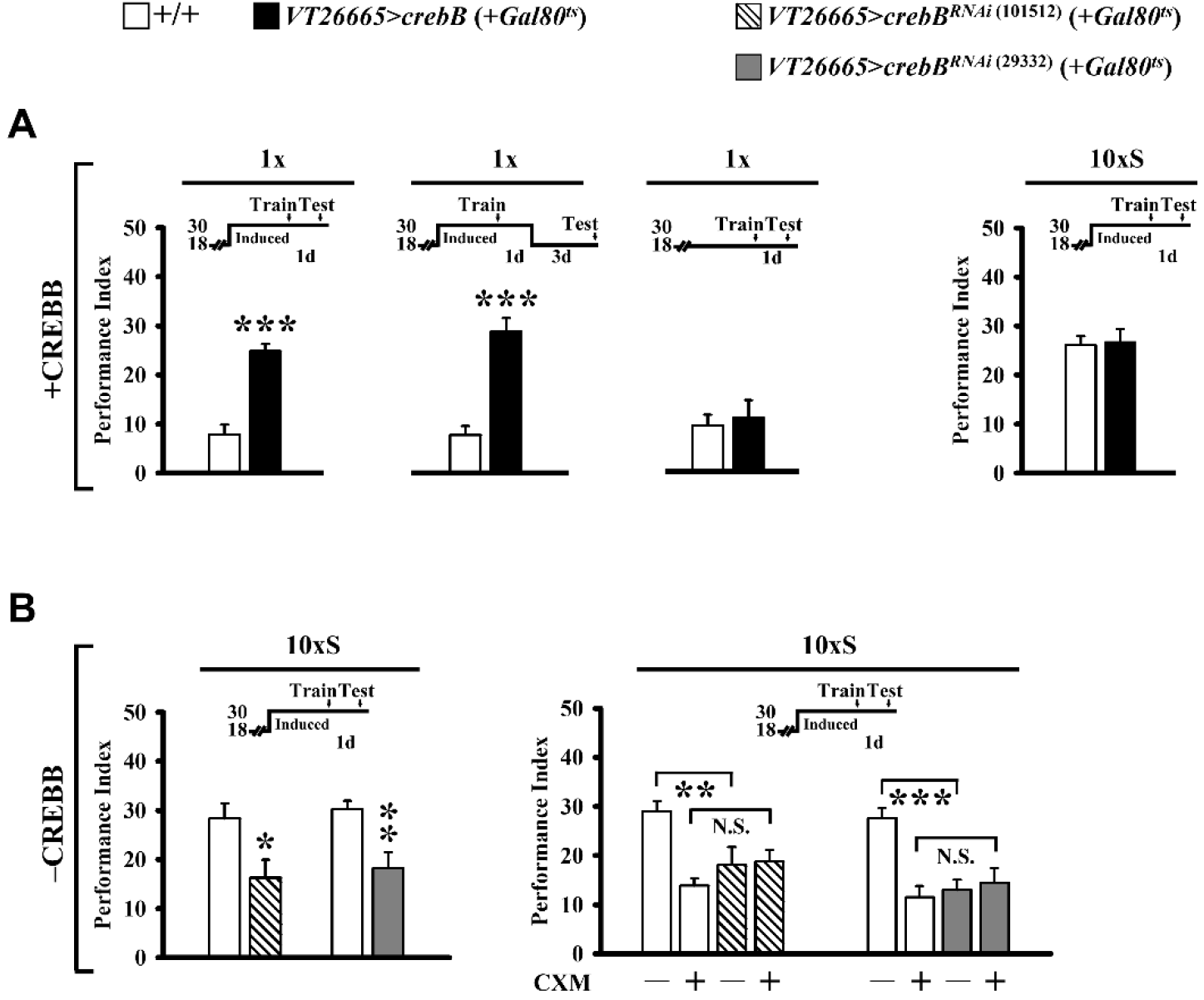
Modulation of CREBB protein in early α/β neurons bi-directionally regulates LTM formation. (**A**) Overexpressing CREBB proteins in early α/β neurons enhances 1-day memory after only 1x training (left) and lasts at least four days (center). *Gal4*-targeted *crebB* overexpression is induced at the restrictive temperature for *tub-Gal80^ts^* (30 °C) from five days before training until testing. Memory is unaffected in these flies held at the permissive temperature for *tub-Gal80^ts^* (18 °C) after 1x training (right). One-day memory is also unaffected in these flies held at 30 °C after 10xS training. (**B**) By contrast, adult-stage specific RNAi down-regulation of CREBB proteins in early α/β neurons *(with two independent RNAi constructs) impairs* 1-day memory after 10xS training (left) (also see figure supplement 6). The same LTM is similarly blocked by systemic protein synthesis inhibition induced by CXM feeding (right).

### Spaced training induces *crebB* transcription

We next generated a *crebB* promoter-driven *Gal4* transgene containing an 11-kb 5′ genomic sequence just upstream of CREBB (see Methods) (Yin et. al., 1995b). This *crebB-Gal4* drives GFP expression in most glia cells and brain neurons, including most MB neurons, though higher levels of expression can be seen in α/β compared to α′/β′ or γ neurons (Figure 9A). By photo converting pre-existing green KAEDE to red prior to training (Chen et. al., 2012), we measured significantly more newly synthesized *crebB-Gal4* green KAEDE in the MB α-lobe during 24-h intervals after 5xS or 10xS training, but not after 1x training or 10xM training in comparison with naïve control flies (Figure 9B-C). This training-induced increase in *crebB* KAEDE appeared specific to the MB neurons because spaced training did not significantly change the levels of new *crebB* KAEDE in ellipsoid body (EB) or glia (Figure 9B, right). These results demonstrate that multiple sessions of spaced training increases CREBB expression in early α/β neurons. Our finding that inhibition of protein synthesis in early α/β neurons enhanced LTM formation (Figure 1-3) suggests that *crebB* gene products function to repress protein synthesis.

**Figure 9.**
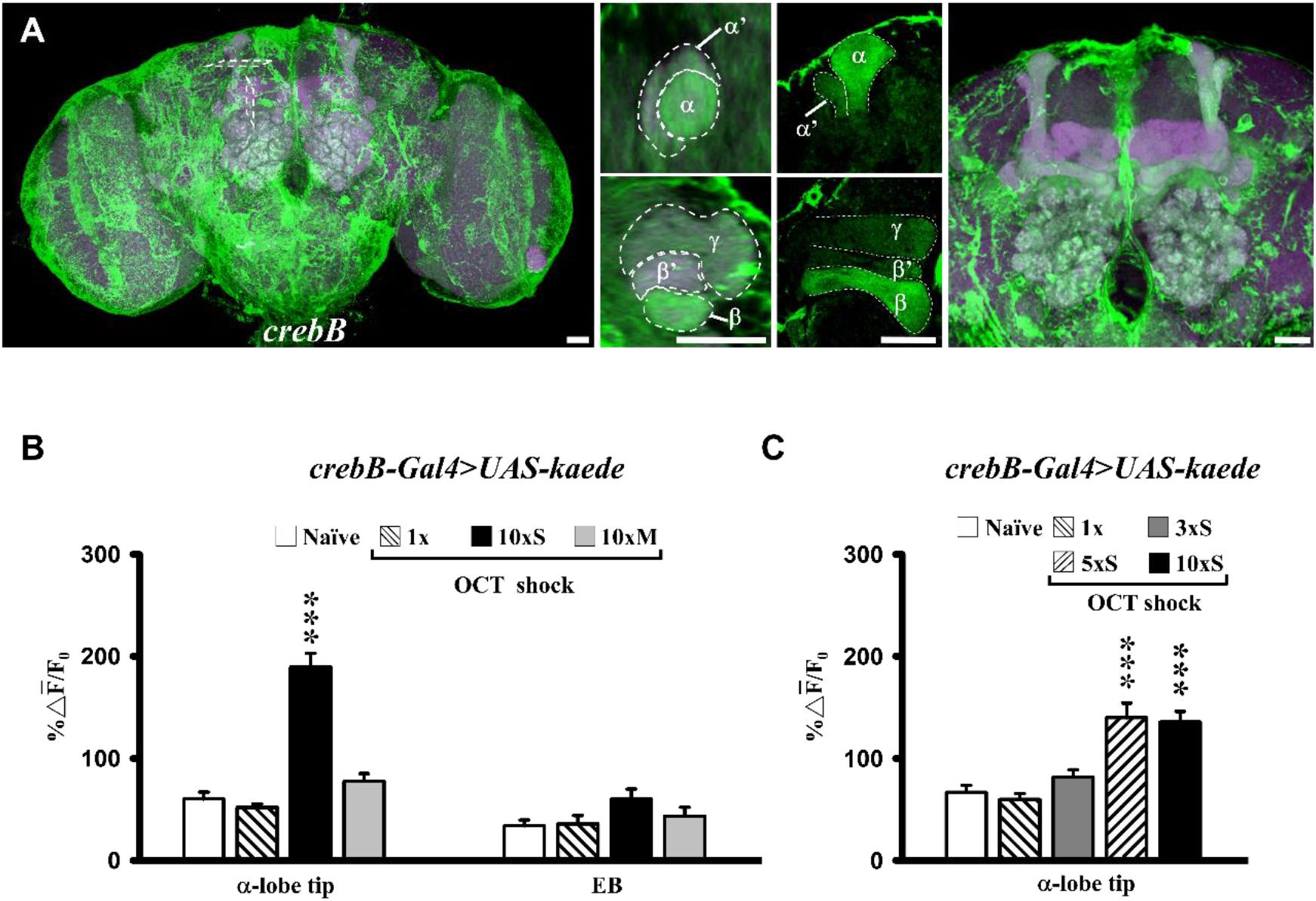
Spaced training activates *crebB* transcription. (**A**) CREBB expression visualized in dissected brains with *crebB-Gal4* driven *UAS-mCD8::GFP* (green), counterstained with DLG-antibody immunostaining (magenta), and viewed under a confocal microscope. Cross sections of vertical and horizontal MB lobes (center) show more prominent expression in α/β neurons than in α′/β′ and γ neurons (labeled). Scale bar = 10 μm. (**B-C**) Promotor activation of *crebB* 24 h after training reported by *de novo* Kaede synthesis, estimated by the ratio of new (green, 488 nm) and preexisting (red, 561 nm) protein 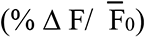. For each brain, single optical slices through the MB α-lobe tip or ellipsoid body (EB) were imaged under identical conditions. (**B**) Spaced training stimulates *crebB* activity preferentially in the α-lobe, in comparison with EB controls. *(B)* A minimum of 5xS training cycles are necessary to observe Kaede synthesis reflecting *crebB* activity. Bars represent mean ± SE, *n* ≥ 8.

### Schnurri (Shn) regulates CREBB-dependent LTM formation

How does spaced training induce CREBB expression? To address this question, we sought to identify positive regulators of *crebB* transcription during LTM formation. Yeast two-hybrid and chromatin immunoprecipitation experiments previously revealed several such candidates that bind in the *crebB* promoter region (data not shown). Prominent among these were (1) CREB family protein CREBA, a leucine-zipper transcription factor (Smolik et al., 1992) and (2) Shn, a zinc finger C2H2 transcription factor encoded by the *shn* gene (Marty et al., 2000) which also was identified in a transposon mutagenesis screen for impairment of 1-day memory after 10xS as the *umnitza* mutant (Dubnau et al., 2003) (described above, see figure supplement 1).

Together, these findings implicated CREBA and Shn as candidate regulators of LTM formation through transcriptional activation of *crebB*. Interestingly, we inducibly overexpressed each in early α/β neurons and found enhanced 1-day memory after 1x or 3xS training with this manipulation of the *shn*^+^ transgene but not *crebA*^+^ (Figure 10A; figure supplement 7A-B). We also observed strongly elevated CREBB protein expression in transgenic *shn^+^* flies, but not in transgenic *rut^+^* or *Pka^act1^* flies (figure supplement 8). Moreover, inducible RNAi knockdowns of *shn* but not *crebA* impaired 1-day memory after 10xS (Figure 10B; figure supplement 7C), and further impairment was not seen with systemic protein synthesis inhibition after CXM feeding (Figure 10B, right). Memory after 10xS was fully rescued in *shn* knockdown flies by *crebB* co-expression (Figure 10C) and was enhanced relative to controls after 1x (Figure 10D). Taken together, these results show that CREBB expression in early α/β neurons in response to spaced training is Shn-dependent.

**Figure 10.**
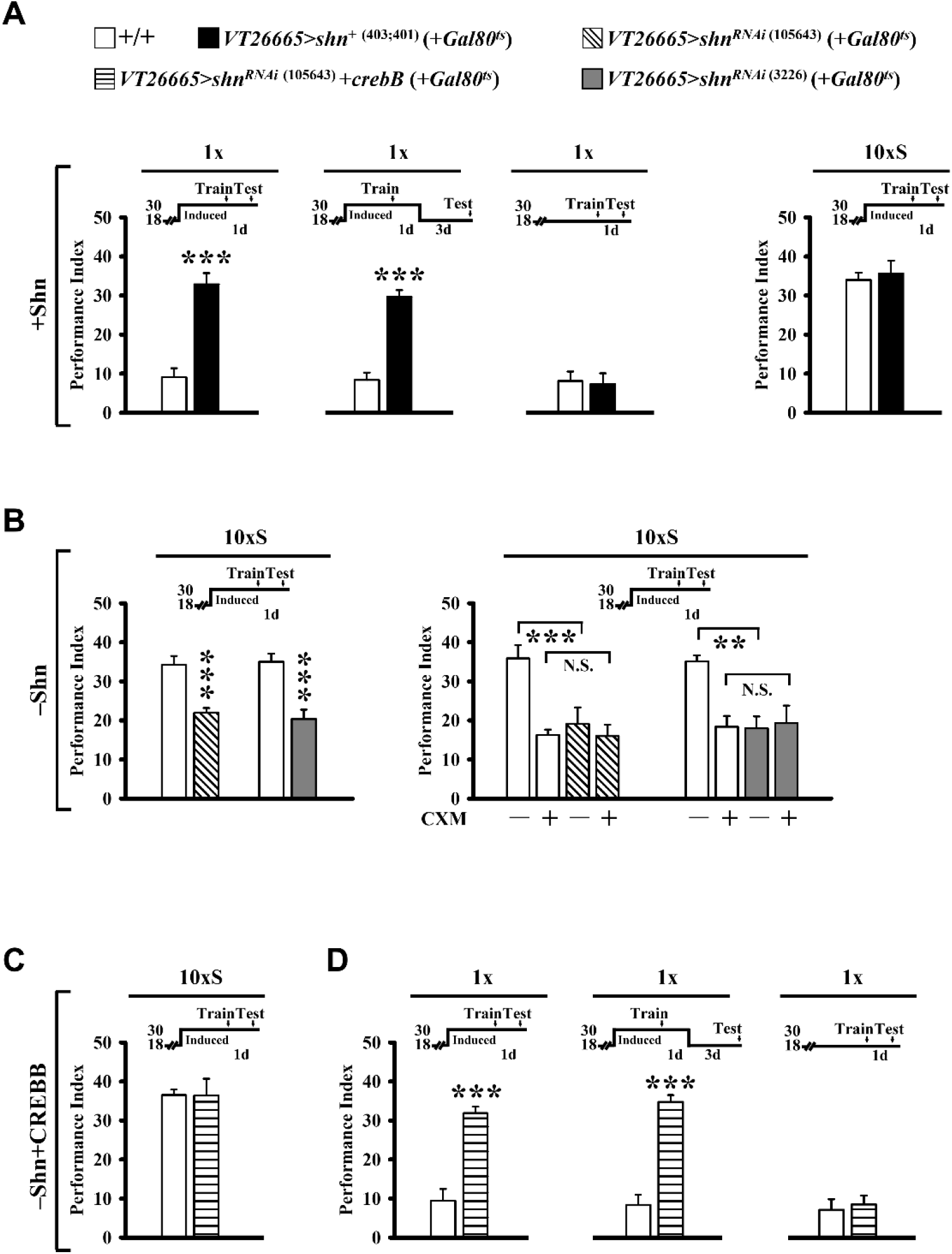
Shn in early α/β neurons regulates CREBB dependent LTM formation. (**A**) Overexpressing Shn proteins in early α/β neurons enhances 1-day memory after only 1x training (left) and lasts at least four days ((left center). *Gal4*-targeted *shn*^+^ overexpression is induced at the restrictive temperature for *tub-Gal80^ts^* (30 °C) from five days before training until testing. Memory is unaffected in these flies held at the permissive temperature for *tub-Gal80^ts^* (18 °C) after 1x training (right center) (also see figure supplement 7). One-day memory is also unaffected in these flies held at 30 °C after 10xS training (right). (**B**) In contrast, adult-stage specific RNAi down-regulation of Shn in early α/β neurons *(with two independent RNAi constructs) impairs* 1-day memory after 10xS training (left) (also see figure supplement 6). The same LTM is similarly blocked by systemic protein synthesis inhibition induced by CXM feeding (right). (**C**) Down-regulating Shn but co-overexpressing CREBB in early α/β neurons does not impair 1-day memory after 10xS training. (**D**) One-day memory is enhanced after only 1x training (left) and lasts at least four days (center). Memory is unaffected in flies held at the permissive temperature for *tub-Gal80^ts^* (18 °C) after 1x training (right).

## Discussion

Our data suggest that MB neurons provide a compelling cellular gating mechanism for LTM formation. A single training session is sufficient to increase early α/β neuronal excitability, the output from which produces a downstream inhibitory effect on LTM formation. After spaced training, cAMP signaling regulates neural excitability and/or Shn increases CREBB expression, the net effects of which we suggest then represses further protein synthesis, thereby reducing early α/β output and relieving the inhibitory effect on LTM formation. Remarkably, our observations emerged from a screen of enhancer trap memory mutants using Ricin^CS^ protein synthesis inhibition (figure supplement 1). 1-day memory after 10xS was impaired in nine lines, eight of which showed expression in both MB & DAL neurons. Curiously, another seven lines were not impaired in 1-day memory after 10xS training – but they, too, showed enhancer expression patterns in MB & DAL neurons. We hypothesized that blocking protein synthesis in DAL neurons impaired LTM but doing so in (some) MB neurons might actually enhance LTM, negating the inhibitory effects in DAL neurons.

We tested this idea by maintaining Ricin^CS^ expression in MB while blocking Ricin^CS^ expression outside of MB using *cry-Gal80* (Figure 1). Surprisingly, LTM now was enhanced in all seven of these enhancer trap lines (Figure 1). We then identified early α/β as the subset of MB neurons responsible for this enhancing effect (Figure 2). Inhibition protein synthesis in early α/β neurons during the first 6 h, or blocking synaptic transmission from early α/β neurons during the first 8 h after training was sufficient to enhance LTM (Figure 3 and 4). Increasing excitability of early α/β neurons impaired LTM, but decreasing excitability again enhanced LTM (Figure 5). We next asked whether these neural excitability-dependent effects were also cAMP dependent. RNAi mediated knockdown of Rutabaga or PKA in early α/β impaired LTM, while overexpression of a *rut^+^* or *Pka^act1^* transgene enhanced LTM (Figure 6 and 7). CREBB expression is suggest to be synergistically and post-transcriptionally regulated by protein kinases responding to cAMP signaling (15) and accordingly, our RNAi mediated knockdown of CREBB in early α/β impaired LTM, while overexpression of a *crebB* transgene enhanced LTM (Figure 8). Finally, using a *crebB* promoter driven *Gal4* transgene, we show that CREBB transcription increases after 5xS or 10xS spaced training but not after 1x training (Figure 9). Thus, spaced training-dependent expression of CREBB repressor proteins in early α/β neurons blocks this inhibitory output from early α/β neurons, thereby allowing LTM formation (downstream) to proceed.

An enhancing role associated with Shn-induced expression of CREBB repressor is a novel aspect of this LTM gating mechanism (Figure 10 and figure supplement 8). Previous reports have claimed that chronic expression of a CREBB repressor or RNAi transgenes in all α/β neurons impaired 1-day memory after spaced training (Yu et al., 2006; Lee et al., 2018). Chen et al., (2012) documented, however, that these chronic disruptions of CREBB produced developmental abnormalities in MB structure. In contrast, acute induced expression of active Ricin^CS^ or CREBB repressor only in adult α/β neurons did not impair 1-day memory after spaced training (and did not produce structural defects). Using a different inducible system (MB247-Switch) to acutely expresses CREBB in γ and α/β neurons, Hirano et al., (2016) showed a mild impairment of 1-day memory after spaced training. More interestingly, they used various molecular genetic tools to show that interactions among CREBB, CREB Binding Protein (CBP) and CREB Regulated Transcription Coactivator (CRTC) in MB clearly were involved in LTM formation or maintenance, respectively. Using the same inducible gene switch tool, Miyashita et al., (2018) showed a fascinating positive regulatory loop between Fos and CREBB in MB during LTM formation – but they did not show behavioral data pertaining to manipulation of CREBB *per se*- and they did not restrict their experiments to early α/β neurons.

A recent study that features *cyclic AMP-response element* (CRE)*-*driven transgenes is pertinent to this report. Zhang et al., (2015) expressed a *CRE*-*luciferase* transgene in different subpopulations of MB neurons and then monitored luciferase activity in live flies at various times after spaced training. Immediately after spaced training, they showed in some cases luciferase expression decreased (*OK107* expressing in all MB neurons; *c739* expressing in all α/β neurons; *1471* expressing in γ neurons), in others expression increased (*c747* and *c772* expressing variably in all MB neurons) or in some no changes were detected (*c320* expressing variably in γ α′/β′ and α/β subpopulation, *17d* expressing primarily in late α/β and in early α/β neurons). Indeed, these authors point out that, because *CRE*-*luciferase* was expressed in more than one subpopulation of MB neurons, only net effects of CREB function could be quantified. Obviously, such a conclusion must be drawn from any behavioral data collected after CREBB manipulations in multiple subpopulations of MB neurons. Our study provides a dramatic example of this point. By restricting our manipulation only to the early α/β neurons and only in adult stage animals, we show that acute overexpression or knockdown of CREBB enhances or impairs LTM formation, respectively (Figure and that spaced training serves to increase the expression of CREBB in these neurons (Figure 9).

Of particular relevance to our future studies is the curious discovery that output from early α/β neurons specifically *inhibits* LTM formation. We find no evidence of inhibitory transmitter (*i.e.,* GABA) synthesis or signaling in early α/β neurons, however, and others have suggested that memory-relevant MB output synapses are cholinergic (Barnstedt et al.,2016). Thus, we presume that inhibition of LTM lies somewhere downstream in the memory circuit. Furthermore, we note that ARM appears to involve α/β neurons (Lee et al.,2011; Knapek et al.,2011; Scholz-Kornehl and Schwärzel, 2016; Kotoula et al., 2017; Shyu et al., 2019) and to inhibit LTM formation (Isabel et al., 2004; Placais et al., 2012). Thus, a molecular link between ARM and LTM may reside in early α/β neurons.

More generally, our results underscore the need to study behavior-genetic relations in each of the seven MB neuronal subpopulations (Aso et al., 2014) separately before drawing firm conclusions about a role for MB in specific memory phases or in the dynamics of a larger memory circuit involving neurons intrinsic and extrinsic to MB. With the more complex circuitries in vertebrate animal models, such deconstruction of memory formation into specific neuronal subtypes will be even more critical and enlightening.

## Materials and Methods

A collection of *Drosophila P-Gal4* transposon insertions were previously selected in an enhancer trap mutagenesis screen for long-term memory phenotypes (Dubnau et al., 2003). The resultant Gal4 expression patterns in seven of these mutants were leveraged to drive and temporally control cold-sensitive Ricin^CS^ activity to block protein synthesis in the identified neuron subsets. In addition, we spatially restricted Ricin^CS^ activity by inhibiting Gal4 with MB or DAL neuron-specific expression of Gal80. We used an automated olfactory aversive learning task (Tully et al., 1994) and assessed LTM after blocking protein synthesis, inhibiting consolidation in these temporally and spatially restricted domains to identify the subsets of neurons critical for this task. Blocking transmission from these neurons with Gal4-targetted temperature-sensitive Dynamin^ts^ after training was used to test the implicated roles of these neurons in LTM consolidation (Dubnau et al., 2001; McGuire et al., 2001). Spatial and temporal regulation of K^+^ and Na^+^ channel activity with transgene overexpression and RNAi knockdown within these neurons was used to assess the downstream impacts of signaling valence on LTM. Similarly, restricted expression of transgenes was used to examine the training-responsive effects on LTM. We evaluated training-responsive CREBB expression with confocal microscopy using a *Gal4*-targeted UV-sensitive KAEDE reporter system (Chen et al., 2012). In various experiments, flies were fed CXM to provide a systemic level of protein synthesis inhibition. Detailed procedures for all methods are described in the supplementary materials.

### Flies

Fly stocks were maintained on standard corn meal/yeast/agar medium at 25 ± 1 °C or 18 ± 1 °C and 70% relative humidity on a 12:12-h light:dark cycle. All genotypes and sources are listed in table supplement 2.

### Behaviour

Olfactory associative learning was evaluated by training 6- to 7-day-old flies in a T-maze apparatus with a Pavlovian olfactory conditioning procedure (Tully and Quinn, 1985) as described previously (Chen et al., 2012; Pai et al., 2013; Wu et al., 2017). All experiments were conducted in the dark in an environment-controlled room at the required temperatures and 70% relative humidity. The odours used were 3-octanol (OCT) and 4-methylcyclohexanol (MCH). Each experiment consisted of two groups of approximately 100 flies, each of which was conditioned with one of the two odours. Flies were exposed sequentially to two odours that were carried through the training chamber in a current of air (odours were bubbled at 750 ml/min). In a single training session, flies first were exposed for 60 s to the conditioned stimulus (CS ^+^), during which time they received the unconditioned stimulus (US), which consisted of 12 1.5-s pulses of 60 V dc electric shock presented at 5-s interpulse intervals. After the presentation of the CS+ condition, the chamber was flushed with fresh air for 45 s. Then flies were exposed for 60 s to the unpaired CS^−^. To evaluate memory retention immediately after single-session training (acquisition), flies were gently tapped into an elevator-like compartment immediately after training. After 90 s, the flies were transported to the choice point of a T-maze, in which they were exposed to two converging currents of air (one carrying OCT, the other MCH) from opposite arms of the maze. Flies were free to choose between and walk toward the CS^+^ and CS^−^ for 120 s, at which time they were trapped inside the respective arms of the T-maze (by sliding the elevator out of the register), anesthetised, and counted. Flies that chose to avoid the CS^+^ran into the T-maze arm containing the CS^−^, whereas flies that chose to avoid the CS^−^ ran into the arm containing the CS^+^. For each experiment, a performance index (PI_1,2_) = (N_CS−_ – N_CS+_)/(N_CS−_ + N_CS+_) was calculated and averaged over these two complementary experiments, with the final PI = (PI_1_ + PI_2_)/2. Averaging of the two reciprocal scores eliminated any potential biases originating from the machine, naïve odour preferences, or non-associative changes in olfaction. For 24-h memory experiments, flies were subjected to single-session training, training massed together without rest, or training spaced out with 15-min rest intervals. For these training protocols, robotic trainers were used. All genotypes were trained and tested in parallel and rotated among all of the robotic trainers to ensure a balanced experiment. The genetic backgrounds of all fly strains were equilibrated to the “Canton” wild-type background by five or more generations of backcrossing. In *tub-Gal80^ts^* experiments, flies raised at 18 °C were transferred to 30 °C for at least five days before the experiments.

### Pharmacological treatment

To block protein synthesis, flies were fed 35 mM cycloheximide (Sigma) in 5% glucose 1 day before training until immediately before the test (Tully et al., 1994).

### *crebB* promoter construct

To engineer the *crebB* promoter construct, polymerase chain reaction (PCR) was performed using genomic DNA from the wild-type *Canton-S w^1118^* (iso1CJ) fly line as the template together with the forward primer 5′GAAAAGTGCCACCTGCTGCATGTCTACCAACAGTTCGAG 3′ and the reverse primer 5′CCGGATCTGCTAGCGGTTCCAGCTGCTGTCTGTATGAC 3′. A 11.6-kb PCR product was generated and inserted into the pBPGAL4.2Uw-2 vector, was digested with AatII and KpnI using In-Fusion^®^ cloning system (Clontech). The promoter construct was injected into *attP^40^*-containing fly strains to obtain the transgenic fly lines.

### KAEDE measurement

KAEDE is a photoconvertible green fluorescent protein, irreversibly changing its structure to a red fluorescent protein upon ultraviolet irradiation (Ando et al., 2002). Taking advantage of circadian transcription and protein synthesis in the lateral clock neurons, we previously validated *de novo* KAEDE synthesis in *per-Gal4>UAS-kaede* flies, in which it faithfully reports the cyclic transcriptions of the *period* gene. Feeding cycloheximide also suppressed green KAEDE synthesis, while not affecting the already-converted red KAEDE (Chen et al., 2012). To measure the amount of newly synthesised KAEDE in MB neurons, we used procedures adapted from a previous study (Chen et al., 2012). Briefly, pre-existing KAEDE proteins were photoconverted into red fluorescent proteins by 365–395 nm UV irradiation generated from a 120-W mercury lamp. For behavioural testing, approximately 15–20 flies kept in a clear plastic syringe were directly exposed to UV light at a distance of 5 cm for 1 h. Individual neurons expressing KAEDE were directly visualised through an open window in the fly’s head capsule. Living samples were used because the signal- to-noise ratio of green to red KAEDE is greatly reduced after chemical fixation. KAEDE neurons were located in less than 5 s by a fast pre-scanning of red KAEDE excited by a 561-nm laser, to avoid unnecessary fluorescence quenching of green KAEDE during repeated scanning. A single optical slice through the MB α-lobe tip was imaged at a resolution of 1024×1024 pixels under a confocal microscope with a 40× C-Apochromat water-immersion objective lens (N.A. value 1.2, working distance 220 μm). All brain samples in the experiment were imaged with the same optical settings maximised for green and red KAEDE immediately before and after photoconversion, respectively. In all cases, both green KAEDE (excited by a 488-nm laser) and red KAEDE (excited by a 561-nm laser) were measured. By using the amount of red KAEDE as an internal standard to calibrate individual variation, we calculated the rate of increase in green KAEDE synthesis after photoconversion with the formula (ΔF) = %(Ft_1_ –average Ft_0_)/average Ft_0_, where Ft_1_ and Ft_0_ are the ratios of the averaged intensities of green (G) to red (R) KAEDE (Gt_0_/Rt_0_) immediately after photoconversion (t_0_) and at a later specific time point (t_1_), respectively.

### Spatiotemporal inhibition of protein synthesis

Ricin^CS^, a mutated Ricin A chain, inactivates eukaryotic ribosomes by hydrolytically cleaving the N-glycosidic bond (A4324) of the 28S ribosomal RNA subunit at high temperatures (30°C), but not at low temperatures (18°C) (Endo et al., 1987; Endo and Tsurugi, 1987; Moffat et al., 1992; Allen et al., 2002). We previously validated the spatiotemporal effect of Ricin^CS^ inhibition in the *Drosophila* brain using lateral clock neurons. We found that Ricin^CS^ can effectively inhibit ∼80% of protein synthesis at a permissive temperature (30°C), which is quickly reversed to normal levels after shifting to a restrictive temperature (18°C) (Chen et al., 2012). This suggests a quick restoration of ribosomal synthesis once Ricin^CS^ becomes inactive. While active Ricin^CS^ is a potent cytotoxin for inhibiting protein synthesis, it tends not to be lethal, as Ricin^CS^ eventually inhibits its own synthesis (Chen et al., 2012, Moffat et al., 1992; Allen et al., 2002). In the current experiments, two copies of Ricin^CS^ was used to block protein synthesis. All flies were raised at 18°C to keep Ricin^CS^ inactive. Before or after training at 18°C, the *Gal4> UAS-ricin^CS^*;*UAS-ricin^CS^* flies were transferred to 30°C for 24 h to activate Ricin^CS^, and then shifted back to 18°C for 1 h to inactivate Ricin^CS^ before the experiments. Temporal control of Ricin^CS^ activation is indicated in the figures for the relevant experiments.

### Immunohistochemistry

Brains were dissected in phosphate-buffered saline (PBS), fixed with a commercial microwave oven (2,450 MHz, 1100 Watts) in 4% paraformaldehyde on ice for 60 s three times, and then immersed in 4% paraformaldehyde with 0.25% Triton X-100 for 60 s three times. After being washed in PBS for 10 min at room temperature, brain samples were incubated in PBS containing 2% Triton X-100 (PBS-T) and 10% normal goat serum, and then degassed in a vacuum chamber to expel tracheal air for four cycles (depressurizing to –70 mmHg and then holding for 10 min). Next, brain samples were blocked and penetrated in PBS-T at 4 °C overnight, and then incubated in PBS-T containing (1) 1:40 mouse 4F3 anti-DLG antibody (Developmental Studies Hybridoma Bank, University of Iowa) to label Disc large proteins, and (2) 1:500 mouse anti-CREBB α657 antibody (from Jerry Yin (Tubon et al., 2013)) at 4 °C for 1 day. Samples were subsequently washed in PBS-T three times and incubated in PBS-T containing 1:200 biotinylated goat anti-mouse IgG (Molecular Probes) as the secondary antibody at 25 °C for 1 day. Brain samples were then washed and incubated with 1:500 Alexa Fluor 635 streptavidin (Molecular Probes) at 25 °C for 1 day. Finally, after extensive washing, immunolabeled brain samples were directly cleared for 5 min in *FocusClear*, an aqueous solution that renders biological tissue transparent (Chiang et al., 2001), and mounted between two cover slips separated by a spacer ring with a thickness of ∼200 μm. Sample brains were imaged under a Zeiss LSM 780 or 880 confocal microscope with a 40× C-Apochromat water-immersion objective lens (N.A. value 1.2, working distance 220 μm).

### Statistics

All raw data were analysed parametrically with SigmaPlot 10.0 and SigmaStat 3.5 statistical software. All the data including the behaviour Performance Index (PI) or KAEDE image (ΔF) were evaluated via unpaired *t*-test (two groups) or one-way analysis of variance (ANOVA) (> two groups). Data were evaluated with the Mann-Whitney Rank Sum Test in cases of unequal variances. Data in all figures are presented as the mean ± SE. Experiments were replicated using multiple *Gal4* drivers with equivalent expression patterns, and multiple effector genes and reagents that impact shared cellular functions.

## Acknowledgments

We thank the Bloomington *Drosophila* stock center, *Vienna Drosophila RNAi Center (VDRC)* and Kyoto *Drosophila* Genomics Resource Centers (DGRC) for fly stocks. We also thank the Developmental Studies Hybridoma Bank for the antibodies.

## Author contributions

Conceived the project, analysed the data, and wrote the manuscript: C.C.C., J.S.D., T.T. and A.S.C.

Imaging experiments: H.W.L.

Behavioural experiments: C.C.C. and F.K.L.

Generated *creb2-Gal4* transgenic flies: R.Y.J. and L.C.

## Data and materials availability

All data are available in the main text or the supplementary materials.

---

**Figure supplement 1.**
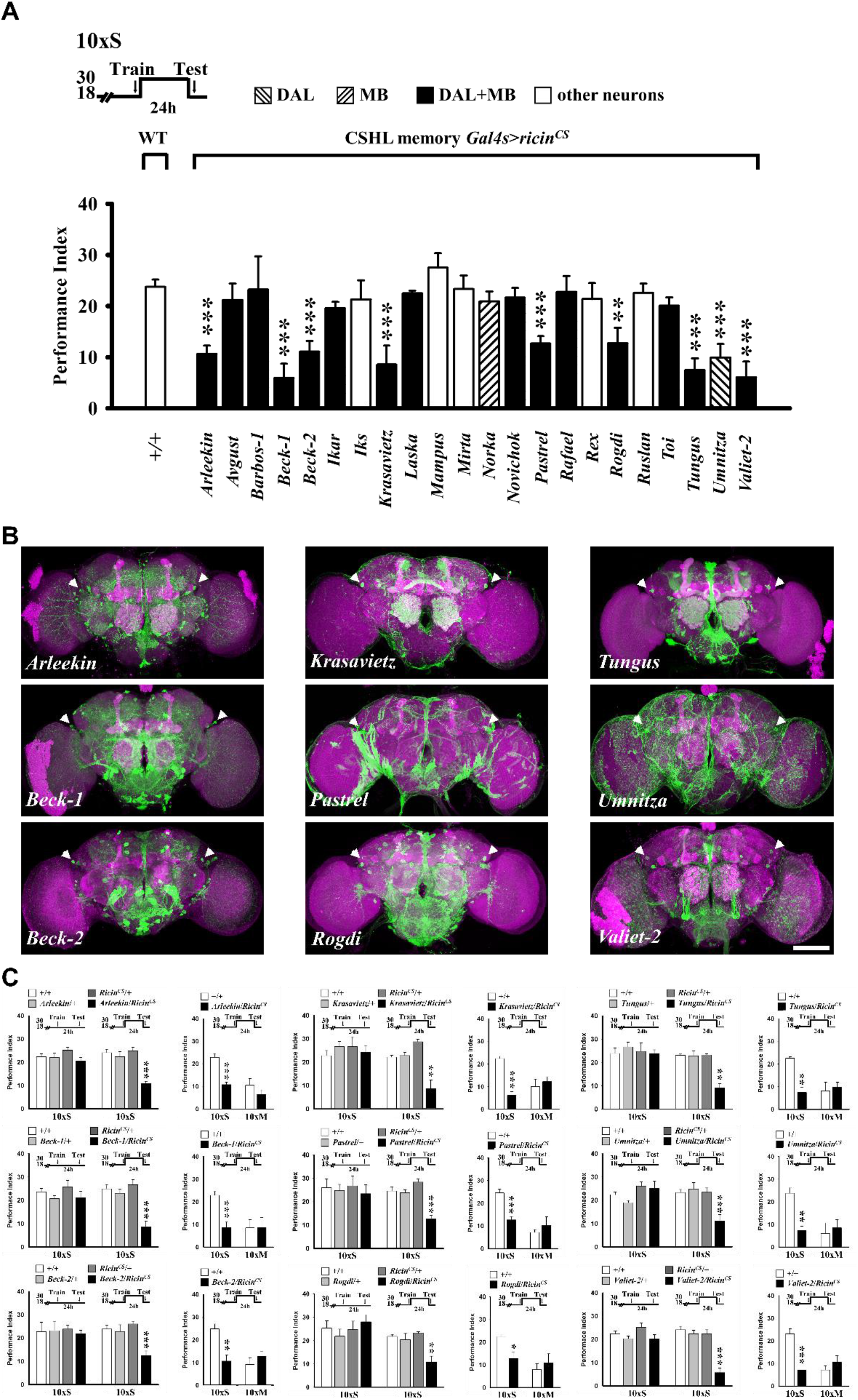
Protein synthesis inhibition in MB and DAL neurons blocks LTM formation. (**A**) Effects of memory circuit *Gal4*-targeted Ricin^CS^ on 1-day memory after ten spaced training cycles (10xS), compared with wild-type (+/+) controls. Cold-sensitive Ricin^CS^ blocks protein synthesis at the permissive temperature (30 °C) between training and testing. Inhibition of protein synthesis in eight of 22 *Gal4-*expressing patterns that include both MB and DAL neurons and one that includes DAL but not MB neurons (Umnitza) blocks 1-day memory. Interestingly, inhibition of protein synthesis in seven other *Gal4-*expressing patterns that include both MB and DAL neurons (related to Figure 1) and one that includes MB but not DAL neurons (Norka) have no net effect on LTM. (**B**) Nine Gal4 expression patterns in which protein synthesis inhibition blocks 1-day memory (**A**). (**C**) We observe no LTM effects after Ricin^CS^ expression in patterns shown above (**B**) at the restrictive temperature (18 °C) or in flies expressing the nine *Gal4* drivers or *ricin^CS^* transgene alone (left). Ricin^CS^ expression in all nine patterns has no effect on 1-day memory after 10xM training (right). In all Extended Data figures, temperature control schedules for behavior experiments are indicated (top). Memory performance indices are calculated as the normalized percent avoidance of shock-paired odor. Bars represent mean ± SE, *n* = 8/bar unless otherwise noted. *, *P* < 0.05; **, *P* < 0.01, ***, *P* < 0.001. All images of dissected brains show Gal4-driven *UAS-mCD8::GFP* (green), counterstained with DLG-antibody immunostaining (magenta), as viewed under a confocal microscope. Scale bar = 50 μm unless otherwise noted. All fly genotypes are listed in table supplement 1.

**Figure supplement 2.**
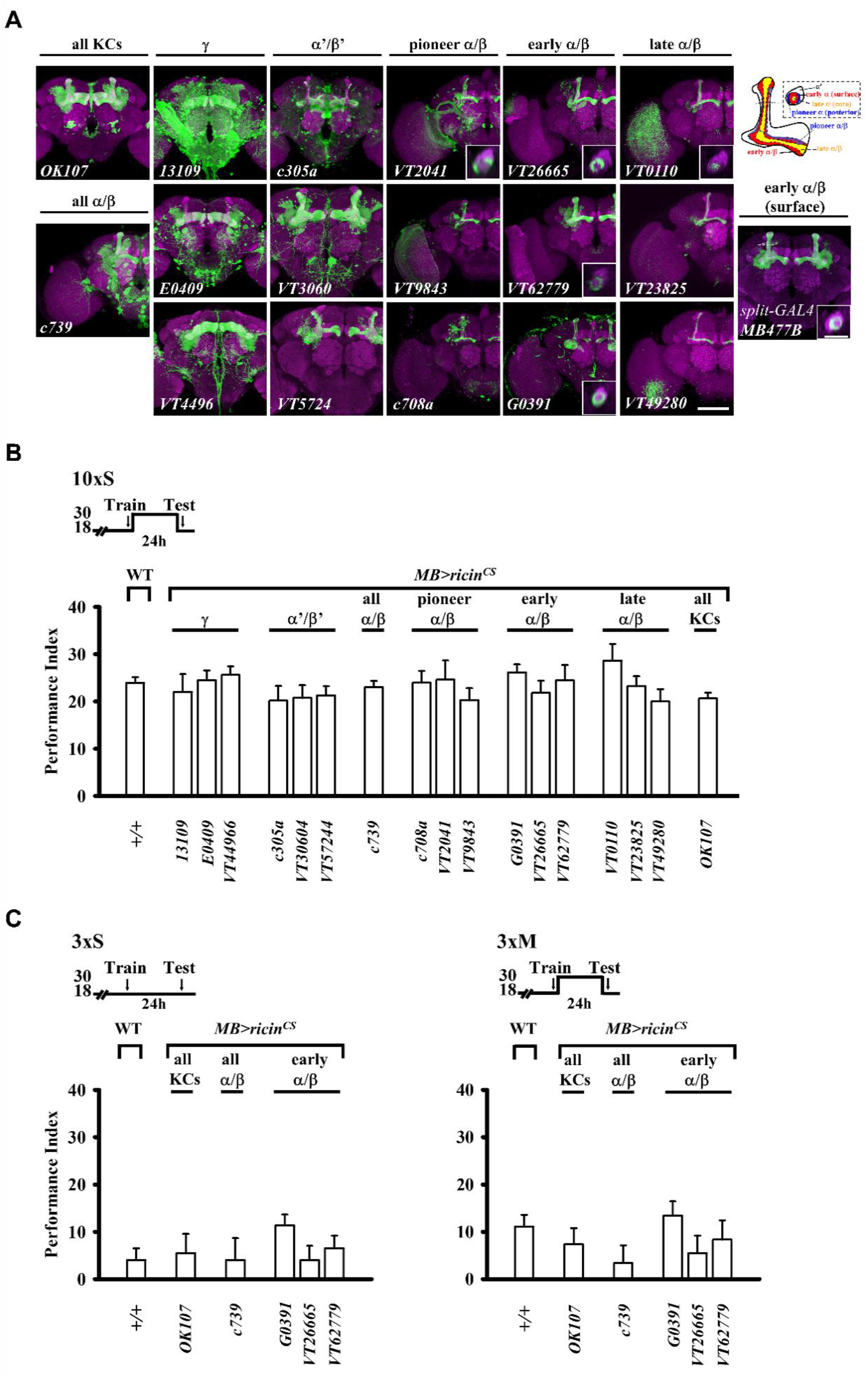
Protein synthesis inhibition in MB neurons does not affect LTM formation after 10xS training. (**A**) *Gal4* expression patterns that delineate five genetically and developmentally distinct MB neuron subtypes. Spatial distributions of three α/β neuron subtypes shown in a schematic representation (right) and in cross section at the vertical lobes (inset). MB *split-Gal4 MB477B* shows specific expression in early α/β (surface) neurons. Scale bar (inset) = 10 μm. (**B**) Effects of MB *Gal4*-targeted Ricin^CS^ on memory compared with wild-type controls (related to Figure 2). Blocking protein synthesis in MB neurons after 10xS training has no effect on LTM (compare with the memory enhancing effects of protein synthesis inhibition in early α/β neurons after 3xS and 1x training, Figure 2). (**C**) Ricin^CS^ expression in MBs neurons has no effect on memory at the restrictive temperature (18 °C) after 3xS training (left), or at the persmissive temperature (30 °C) after 3xM training (right).

**Figure supplement 3.**
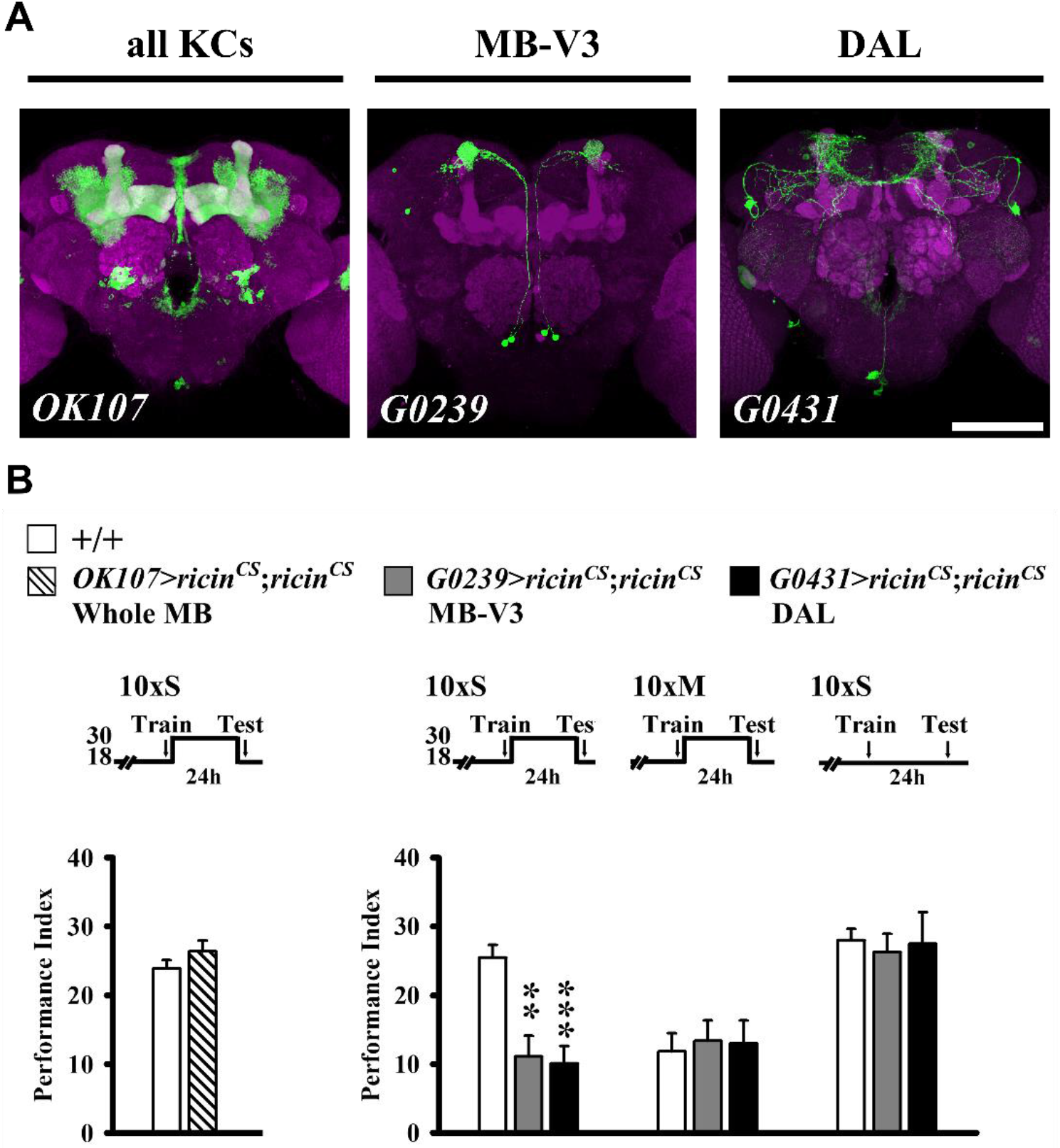
Protein synthesis inhibition in specific extrinsic MB neurons blocks LTM formation. (**A**) *Gal4* expression patterns that delineate intrinsic MB neurons and extrinsic MB-V3 and DAL neurons. (**B**) Blocking protein synthesis in MB neurons with two copies of active Ricin^CS^ has no effect on LTM after 10xS training (left). By contrast, blocking protein synthesis in MB-V3 or DAL neurons inhibits LTM after 10xS training (right), but not after 10xM training or after 10xS training when flies are held at the restrictive temperature (18 °C).

**Figure supplement 4.**
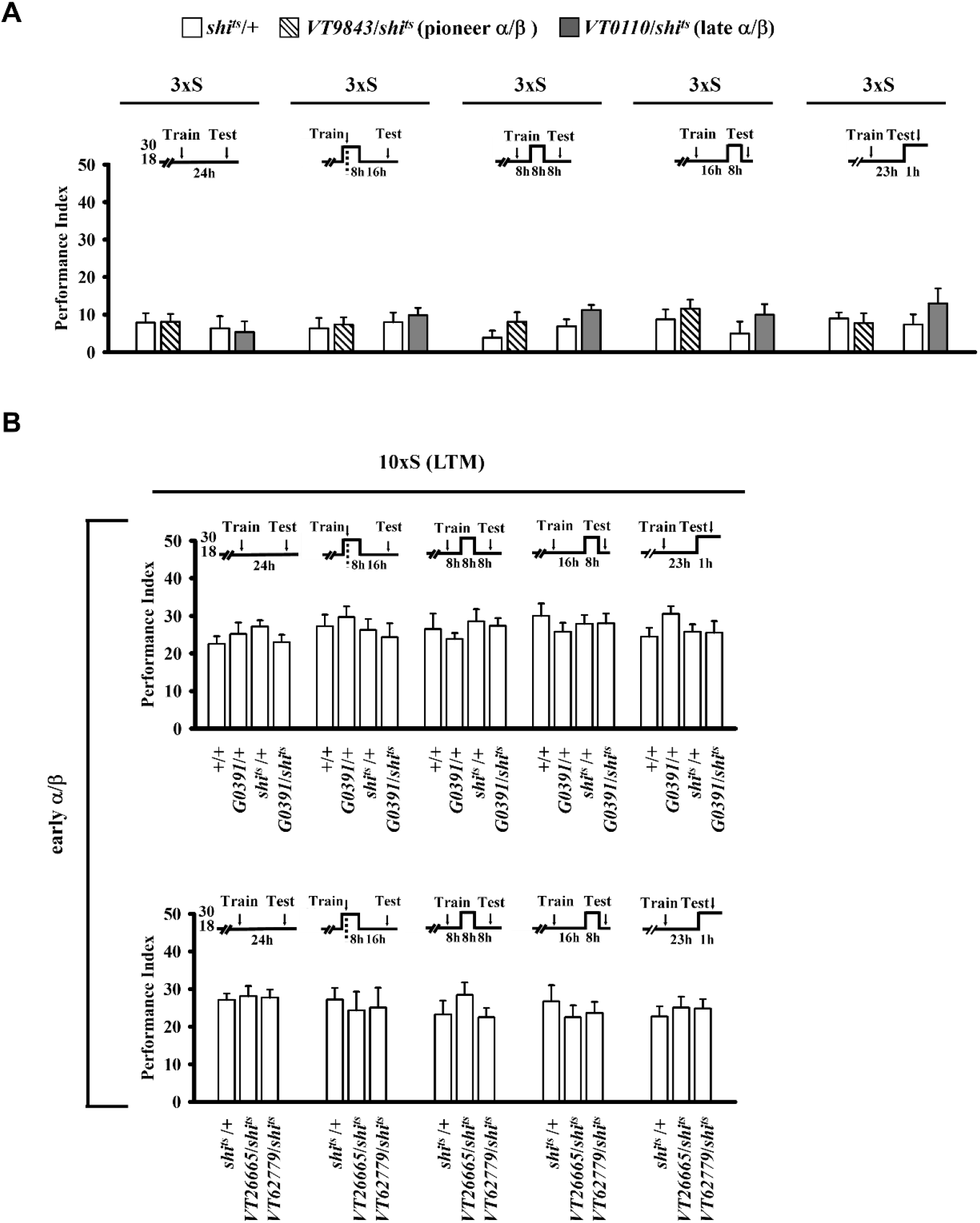
Blocking transmission from MB neurons and effects on LTM formation. Effects of a/b *Gal4*-targeted Dynamin^ts^ (shi^ts^) on 1-day memory. (**A**) Blocking neural transmission from pioneer and late a/b neurons during sequential 8-h time windows after subthreshold 3xS training has no effect on memory. (**B**) Blocking neural transmission from early a/b neurons during sequential 8-h time windows after 10xS training has no effect on memory. By comparison, blocking transmission from early a/b neurons after 3xS training enhances memory (Figure 4).

**Figure supplement 5.**
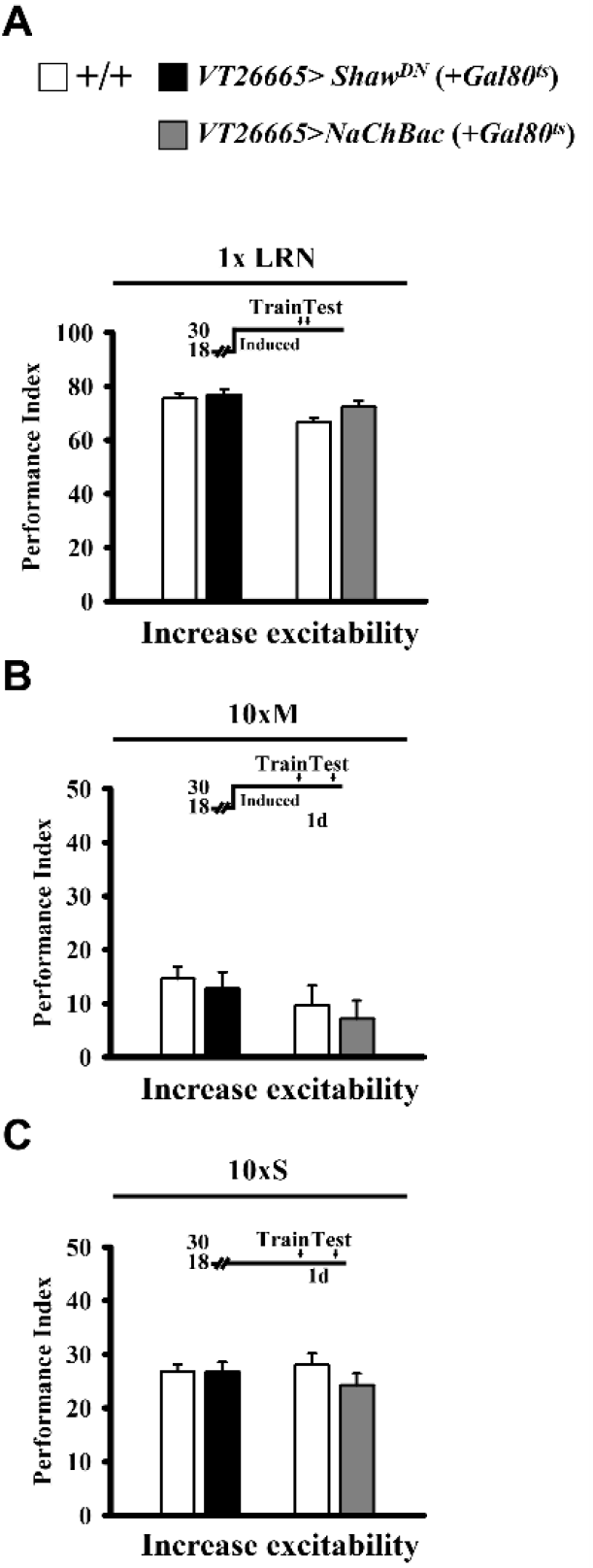
Increasing membrane excitability in early α/β neurons and effects on memory formation. Early a/b *Gal4*-targeted expression of K^+^ and Na^+^ channel proteins is induced at the restrictive temperature for the *tub-Gal80^ts^* inhibitor (30 °C) from five days prior to training until testing. Experimental groups are compared with wild-type (+/+) controls. (**A**) *Shaw^DN^* and *NaChBac* overexpression increase neural activity in early α/β neurons but have no effects on immediate memory after 1x training. (**B**) Similarly, elevated neural activity has no effect on 1-day memory after 10xM training. By comparison, elevated neural activity impairs 1-day memory after 10xS training (Figure 5). (**C**) Channel proteins are not induced in flies maintained at the permissive temperature for the *tub-Gal80^ts^* inhibitor (18 °C) and show no differences in memory after 10xS training.

**Figure supplement 6.**
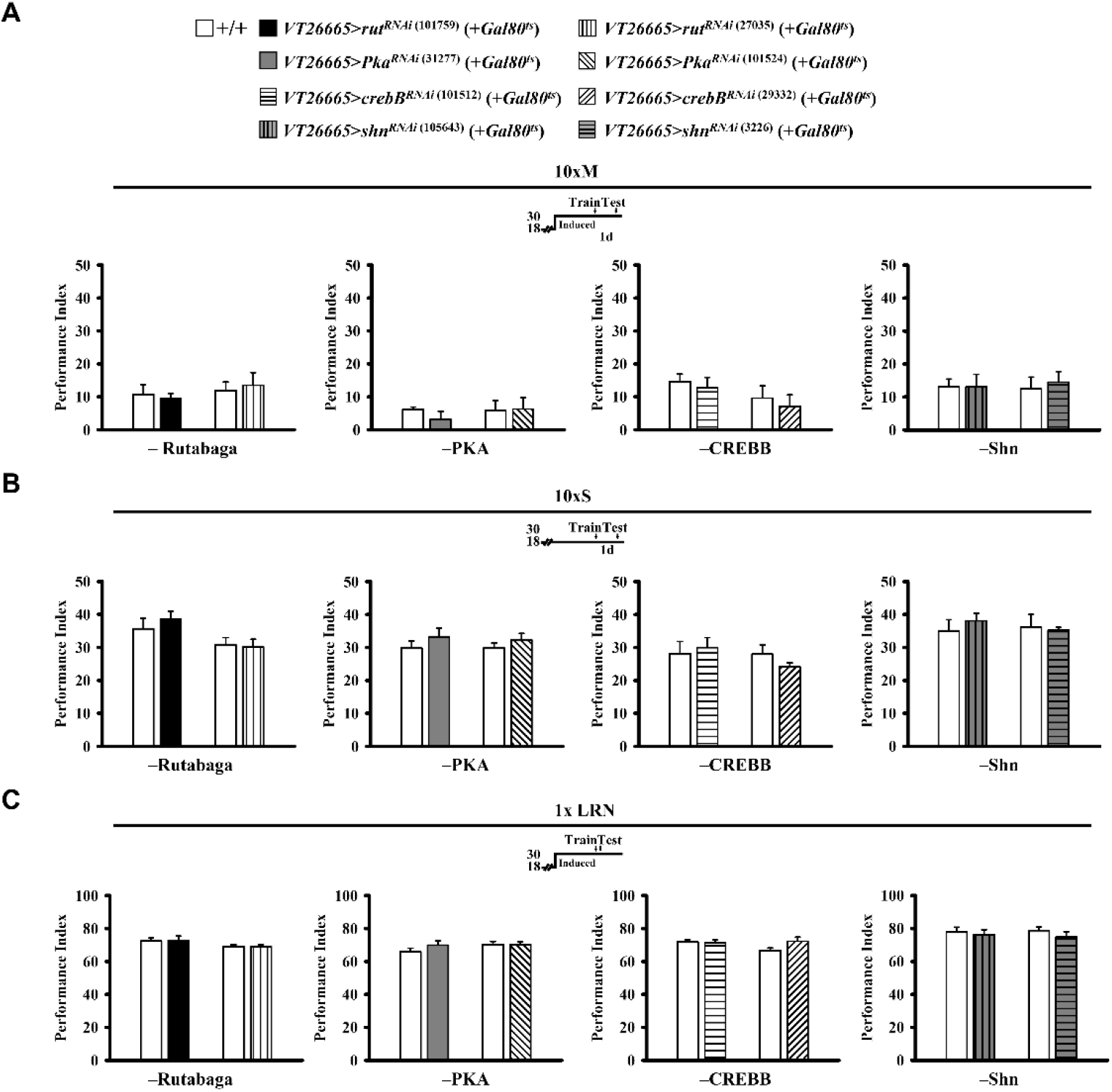
Down-regulation of CREBB, Rutabaga (AC), PKA and Schnurri (Shn) in early α/β neurons and effects on memory formation. (**A**) Adult-stage specific RNAi down-regulation of these proteins (encoded by *rut*, *Pka*, *crebB* and *shn*) in early α/β neurons (two RNAi constructs each) have no effects on 1-day memory after 10xM training. *Gal4*-targeted RNAi specific for each gene is induced at the restrictive temperature for *tub-Gal80^ts^* (30 °C) from five days prior to training until testing. Experimental groups are compared with wild-type (+/+) controls. (**B**) Proteins are not down-regulated in flies maintained at the permissive temperature for the *tub-Gal80^ts^* inhibitor (18 °C) and there are no differences in memory in comparison with control flies after 10xS training. (**C**) Down regulation of all proteins in early α/β neurons has no effect on immediate memory after 1x training. By comparison, overexpressing AC, PKA, CREBB and Shn in early α/β neurons enhances 1-day memory after only 1x training (Figure 6-8 and 10).

**Figure supplement 7.**
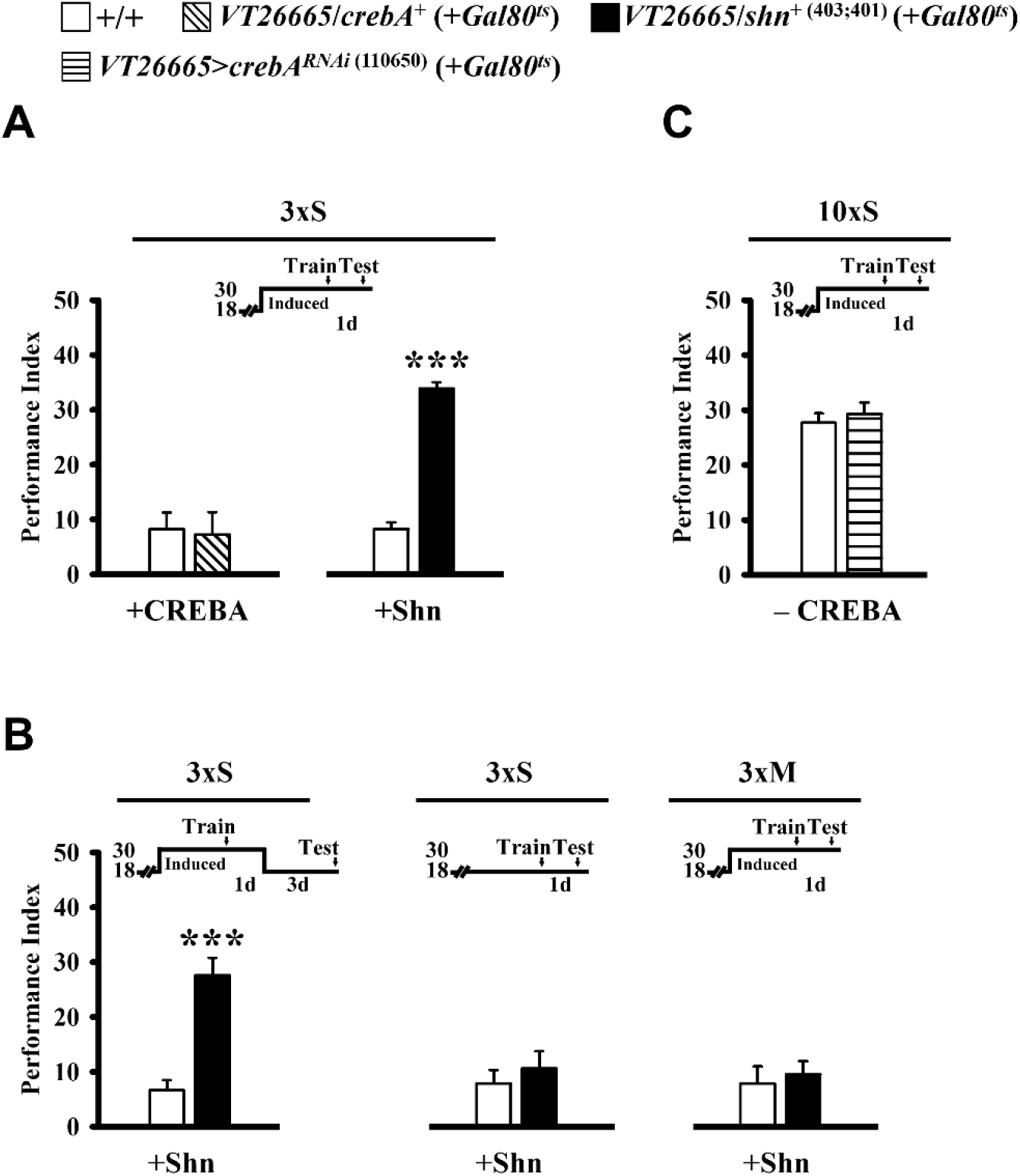
Overexpression of Shn but not CREBA protein in early α/β neurons enhances LTM formation. (**A**) Overexpressing Shn but not CREBA in early α/β neurons enhances 1-day memory after 3xS training. *Gal4*-targeted *crebA* or *shn* overexpression is induced at the restrictive temperature for *tub-Gal80^ts^* (30 °C) from five days prior to training until testing. (**B**) Enhanced memory lasts at least four days (left). Memory is unaffected in these flies held at the permissive temperature for *tub-Gal80^ts^* (18 °C) after 3xS training (center), or at the restrictive temperature (30 °C) after 3xM training (right). (**C**) Adult-stage specific down-regulation of CREBA protein in early α/β neurons *has no effect on* 1-day memory after 10xS training.

**Figure supplement 8.**
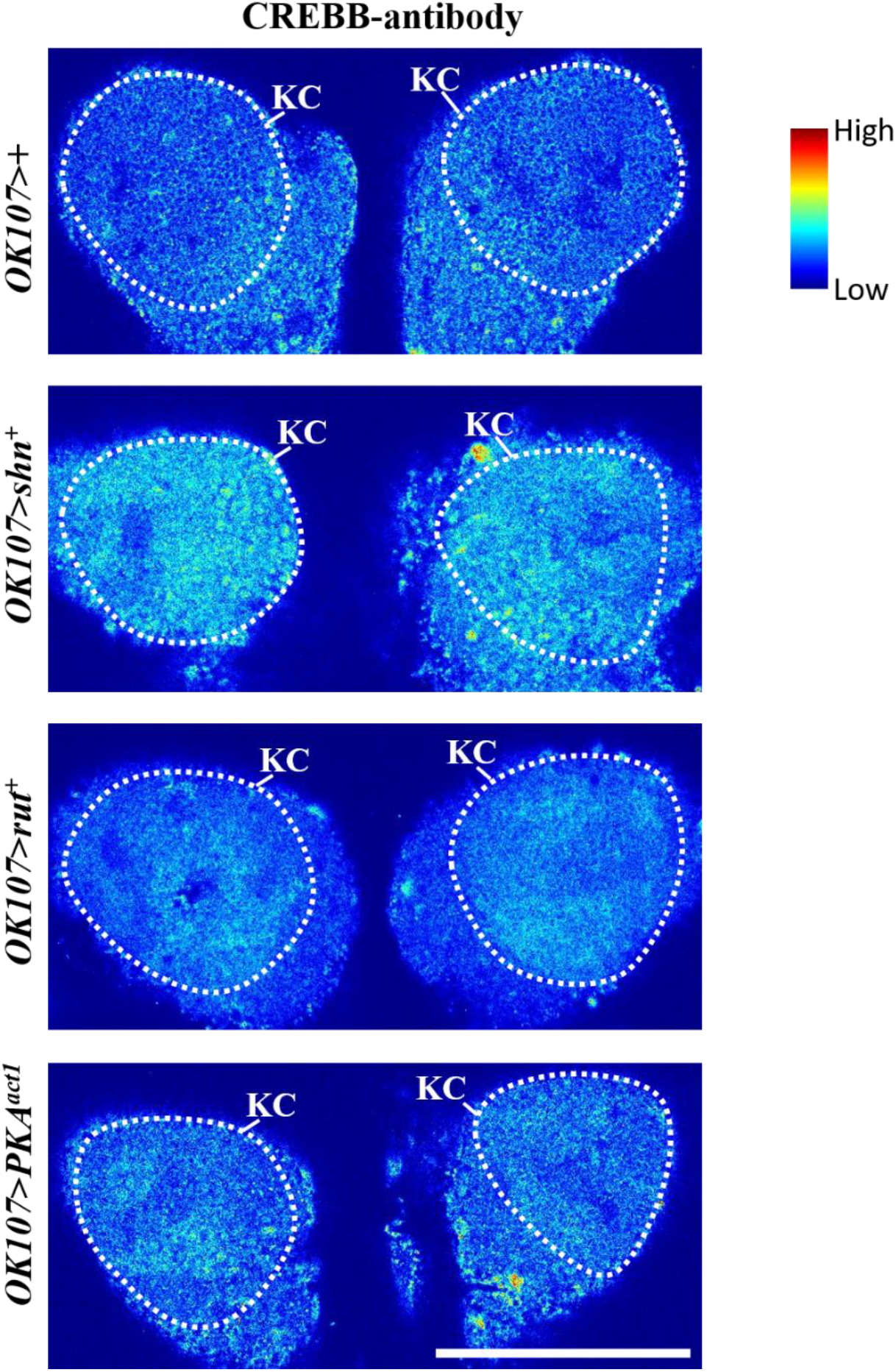
Transgene over-expression and CREBB protein levels in MB Kenyon cells. *Gal4-OK107-*driven *shn* over-expression led to elevated CREBB protein levels (second from top) in comparison with the control (top). Over-expression of *rut* and *PKA* had only minor impact on CREBB expression (second from bottom and bottom, respectively). Confocal images of *Drosophila* MB Kenyon cell bodies (encircled by the white dotted line) show the intensity of CREBB immunostaining in a jet colormap.

**Table supplement 1.**
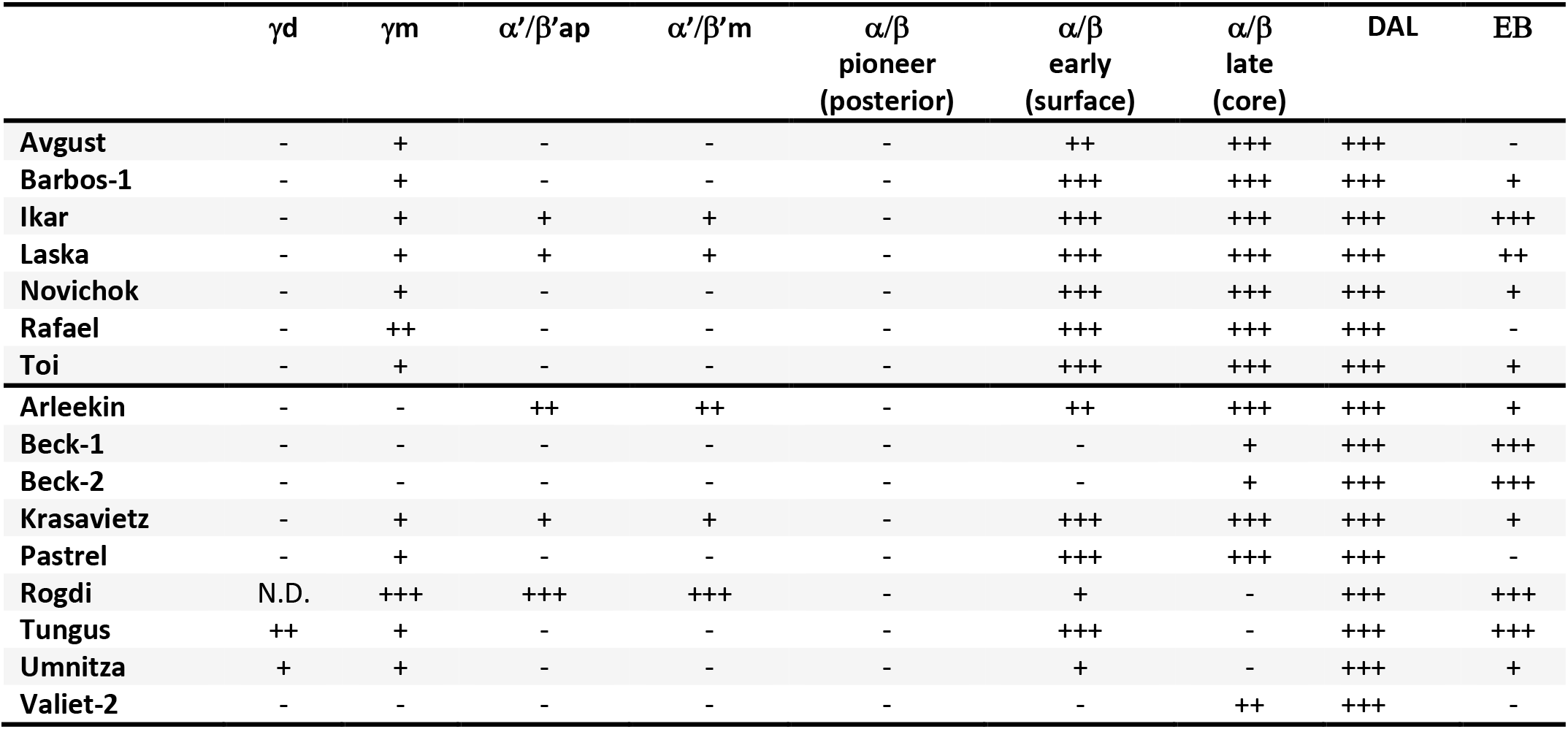
CSHL memory Gal4 patterns. Table shows Gal4 targeted GFP intensity was graded as strong (+++), intermediate (++), weak (+), absence (−) or non-distinguishable (N.D.). DAL, dorsal anterior lateral neuron; EB, ellipsoid body.

**Table supplement 2.**
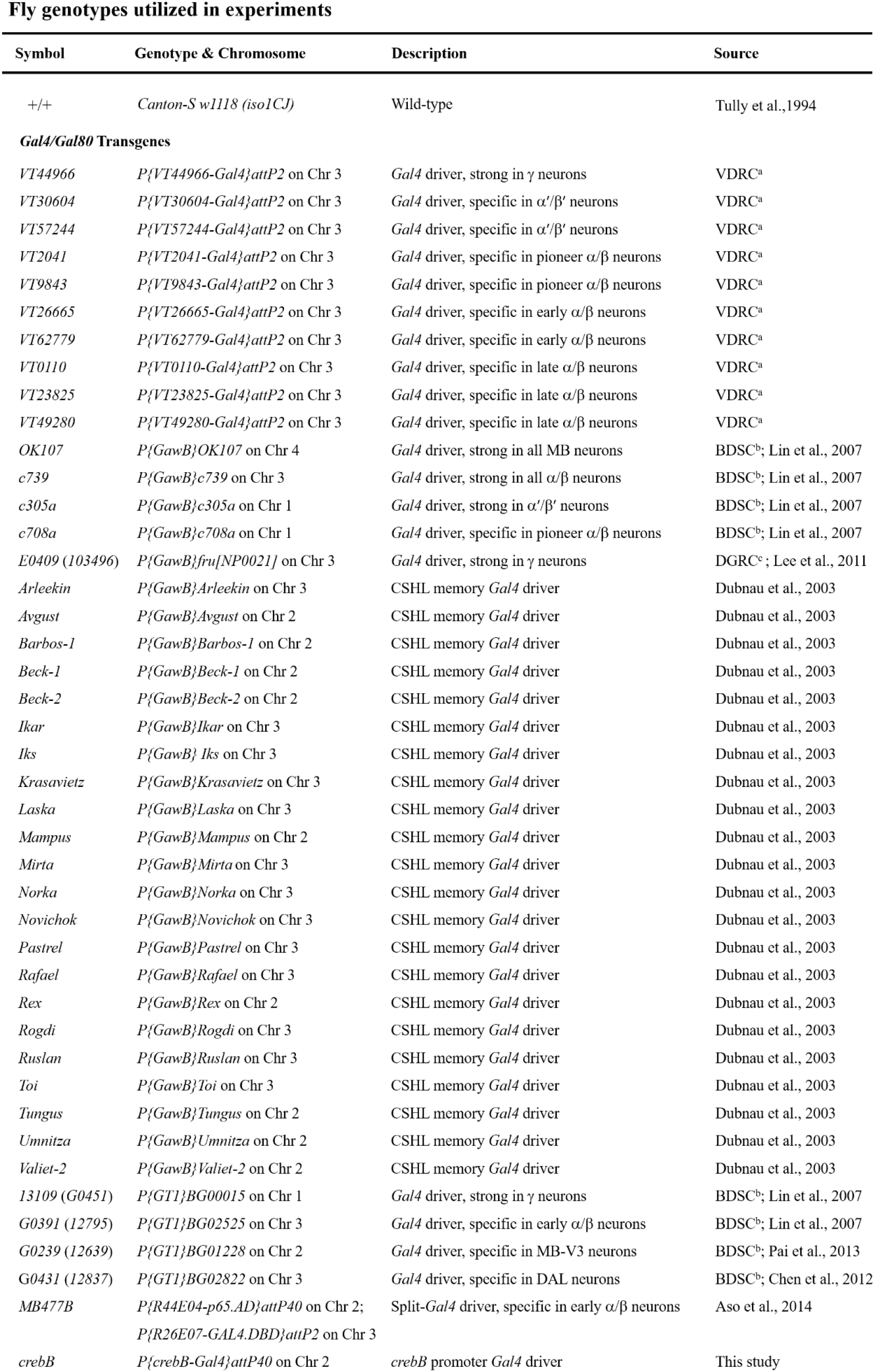
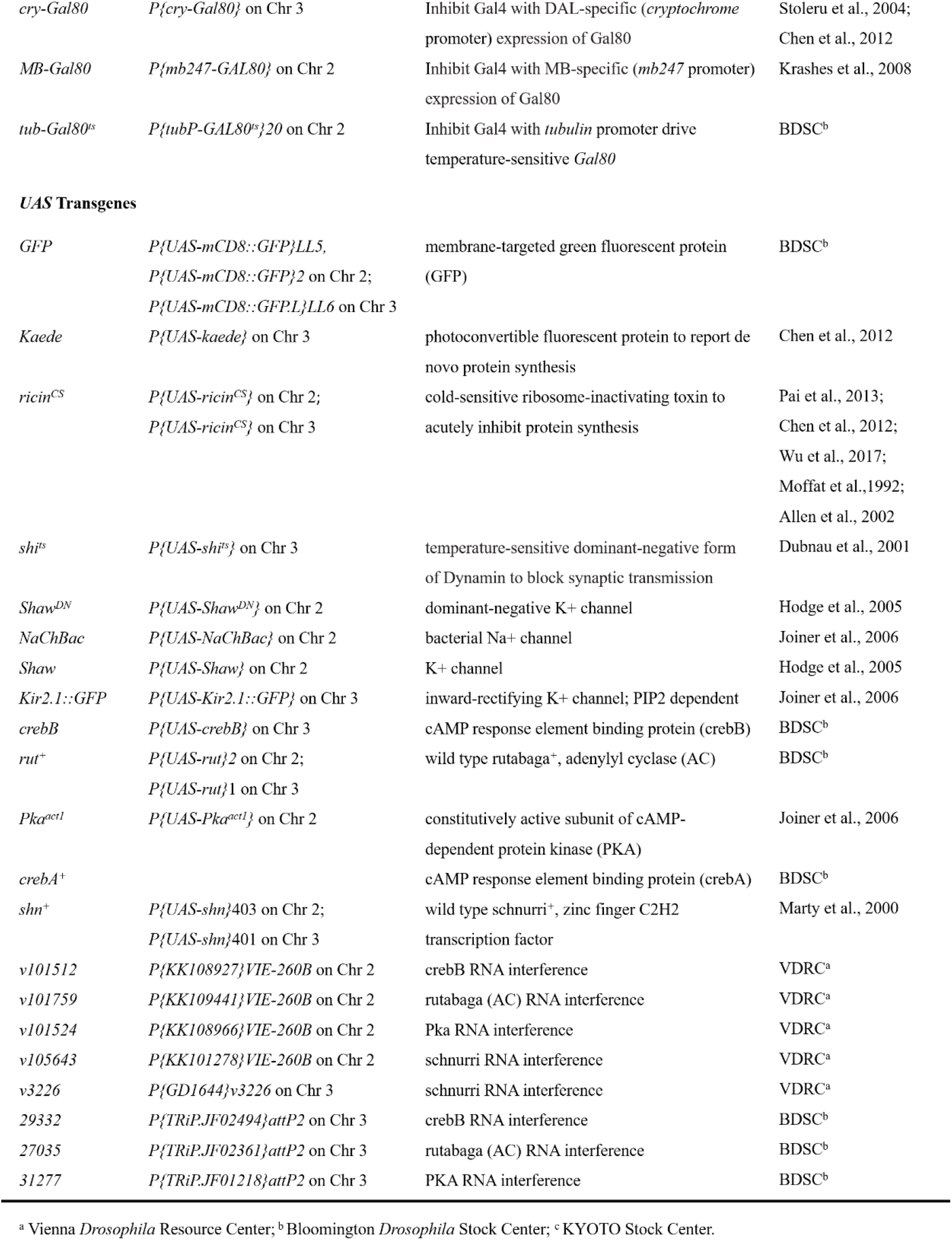
Fly genotypes used in this study. Table shows all fly genotypes, descriptions and sources used in this study, grouped by distinct types of transgenes. References are listed in Methods.

